# Myelinated mechano-nociceptors encode noxious cold and underlie hypersensitivity in pulpitis

**DOI:** 10.64898/2026.09.28.755141

**Authors:** Elizabeth A. Ronan, Shuhao Wan, Sienna K. Perry, Aditi Jha, Akash R. Gandhi, Jingchen Xu, Sabrina T. Wang, Lindsey L. Wu, Kevin P. Pipe, Joshua J. Emrick

## Abstract

The ability to identify dangerous temperatures is critical, but the neural basis for cold sensation remains enigmatic. By combining in vivo functional imaging with tissue-specific spatiotemporal cooling, we unveil myelinated high threshold-mechanonociceptors (HTMRs) as the neural origin of tooth cold sensitivity. Critically, this neural encoding is driven by threshold noxious cold temperature, not cooling rate and remains independent of the cool receptor TRPM8 or pain transducer Nav1.8. Furthermore, while *Bmpr1b*-Cre captures the intradental HTMRs, it further reveals myelinated HTMRs mediate trigeminal cold encoding in soft tissues. Finally, we determine that inflammation potentiates intradental HTMRs, providing functional basis for clinical hypersensitivity in pulpitis. Collectively, our results reveal a new neural mediator of cold sensation and provide insight into painful transformation of cold sensory encoding in inflammation.

## Main Text

Cold temperatures pose severe environmental threats to survival across all levels of physiology, ranging from influencing the rate of essential metabolic processes to damaging tissue irreversibly(*1–3*). In humans and other mammals, low temperature is defined as any sub-baseline body temperature (i.e., ∼30°C for superficial peripheral tissues), further subdivided into innocuous (∼16°C to <30°C) or noxious (≤ 15°C) cold based on both psychophysical measurement and the increasing risk of irreversible bodily harm(*4*, *5*). To detect, perceive, and respond to dangerous thermal conditions, evolution has prioritized mammalian peripheral somatosensory systems that detect local thermal stimuli and relay temperature information to the central nervous system to promote survival. Studies on cold sensation in recent decades have largely centered on characterizing the role of transient receptor potential (TRP) channel TRPM8 in cold sensing, which is a bona fide innocuous cool (and menthol) ion channel receptor that marks distinct somatosensory neurons(*4*, *6–8*). Critically, multiple lines of evidence using in vivo behavioral assays and cellular recordings demonstrate noxious cold sensing persists despite the loss of TRPM8 hinting at alternative neural mediators (*4*, *9–12*). However, detailed studies examining TRPM8-independent mechanisms by which the somatosensory system encodes noxious cold remain limited.

While somatosensory encoding of both heat and mechanical sensation has received major attention(*13–15*), a significant knowledge gap remains in understanding how low temperatures are encoded in the periphery. Single cell sequencing of somatosensory neurons in either the dorsal root ganglia (DRG) or trigeminal ganglia (TG) reveals at least ∼15 molecularly distinct TRPM8-negative subpopulations whose functional tuning has been primarily characterized within cutaneous skin. The functional tuning of these populations has been primarily defined surrounding their encoding of mechanical and/or heat stimuli(*13*, *16–18*). However, comprehensive examination of low temperature encoding mechanisms using this lens, particularly in peripheral tissues other than cutaneous skin, remains lacking.

Progress in understanding somatosensory encoding of cold has been hindered by technical challenges in administering controllable, precise cooling with respect to the anatomical site, target temperature, and cooling rate. Thus, the majority of work examining neural responses to cold stimuli in vivo has relied on widely-available, low-spatial-resolution methods without control and precision (e.g., liquid perfusion)(*19–22*). Recently, emerging tools that utilize the thermoelectric (i.e., Peltier) effect have been applied in neuroscience research to overcome these challenges(*4*). Peltier devices with careful cooling concentrator designs combine cooling power from bigger/multiple thermoelectric components to administer localized cooling to discrete tissues(*23*, *24*). Importantly, Peltier devices utilize electrons as the energy carrier(*4*, *25*), offering rapid control of the temperature setpoint and rate of change by regulating the operating current and thereby enabling careful study of neuronal activity associated with temperature dynamics. Despite this, existing neuroscience research incorporating Peltier devices either primarily focused on heating(*15*) and failed to examine how changes in temperature dynamics influence somatosensory neuron activation(*13*, *26*).

Dental innervation provides a tractable model tissue to explore noxious cold sensation and shed insights into cold encoding in the periphery. Tooth sensation is closely associated with pain, and tooth cold hypersensitivity and is one of the most common clinical complaints related to orofacial pain(*27–29*). We recently demonstrated that tooth sensory innervation is dominated by a discrete population of molecularly specialized trigeminal myelinated high threshold mechano-nociceptors (HTMRs) characterized by their combined expression of the myelinated marker *S100b* and the nociceptive marker *Scn10a*(*30*). Our group established that intradental HTMRs in the intact tooth are specifically tuned to structural enamel damage and that their selective activation drives pain behavior in awake animals, confirming their role as nociceptors(*30*). Critically, tooth inflammation (i.e., pulpitis) is closely associated with heightened cold and mechanical tooth hypersensitivity(*27*, *31*, *32*), hinting that intradental HTMRs could function as the neural mediators of inflammatory tooth pain in mammals.

Here, we combine precision spatiotemporal cooling with a powerful in vivo calcium imaging platform that enables rapid identification and monitoring of intradental sensory innervation (*30*, *33*). Taking inspiration from the design of a cryoanesthesia/analgesia applicator(*34*), we custom-engineer Peltier probes with cooling concentrators designed to deliver high cooling power to miniature tip areas, providing the capability of targeting discrete intraoral and orofacial structures while administering rapid and controllable cooling. We utilize precise control of temperature dynamics to examine intradental HTMR responses to features of cooling and reveal that intradental HTMRs are tuned to a noxious temperature threshold independent of relative temperature change and cooling rate. We confirm that these cold encoding HTMRs do not express the canonical cool sensor *Trpm8,* in alignment with single cell RNA sequencing data(*13*, *16–18*) and our prior multi-plexed in situ hybridization (ISH) studies characterizing somatosensory populations(*35*, *36*). Moreover, genetic knockout of *Scn10a*/Nav1.8 (a putative cold pain transducer) resulted in no deficit in intradental cold responses, further indicating that intradental HTMR cold encoding may rely on noncanonical noxious cold encoding mechanisms. We find that *Bmpr1b*-Cre, which labels a subpopulation of mechanonociceptors(*13*, *37*), marks both intradental HTMRs in addition to a cold-sensitive HTMR population that innervates the lip. We further reveal that in pulpitis, intradental HTMRs display potentiated mechanical and cold responses with no change in spontaneous activity, providing a neural and functional basis for tooth inflammatory hypersensitivity. Taken together, our study sheds insight into how inflammation can transform sensory encoding of cold and mechanical stimuli in teeth and reveals a new neural mediator for cold temperature encoding in the periphery. This work indicates that selective targeting of intradental HTMRs may serve as a useful therapeutic target for inflammatory cold pain in the teeth, and hints at potential conserved function across additional peripheral tissues.

## Results

### Endo-Ice™ refrigerant activates intradental high threshold mechano-nociceptors (HTMRs)

To diagnose pulpal disease, dentists assess the viability and sensitivity of a tooth in response to application of extremely cold refrigerant spray (i.e., Endo-Ice™; -26°C) to the superficial enamel(*38–43*). Patients with tooth pulp ‘health’ describe this experience as evoking an immediately reversible (1-2 second), quick and sharp sensation(*42*, *43*). By contrast, Endo-Ice™ application to a tooth with inner inflammation (i.e., pulpitis) evokes rapid onset, heightened pain with a prolonged duration (seconds to minutes)(*24*, *29*). Inspired by these clinical observations, we sought to first establish whether our previously identified intradental HTMRs directly contribute to these phenomena.

To characterize sensory encoding within intradental HTMRs, we recently established an in vivo calcium imaging platform using a live, anesthetized mouse that enables identification and monitoring of intradental HTMR somas associated with a single tooth, distinguishing them from thousands of superficial TG neurons(*30*, *33*). We previously validated this method in conjunction with retrograde tracing strategies to ensure selective activation of intradental HTMRs(*30*). To characterize intradental HTMRs’ functional tuning to select stimuli, we elected to focus our efforts on intradental HTMRs innervating the ipsilateral 1st mandibular molar (m1), given that the molecular identity of intradental HTMRs remains consistent across individual molars(*35*). This prior molecular characterization established that intradental HTMRs co-express the myelinated marker *S100b* as well as *Scn10a*, which encodes the putative nociceptive marker and voltage gated sodium channel Nav1.8(*30*, *44*). Thus, we elected to evaluate cold-temperature evoked responses in intradental HTMRs using the well-established *Scn10a*-Cre driver line(*44*, *45*) crossed with Cre-dependent GCaMP6(f)(Ai95D) that provides reliable and robust GCaMP expression in intradental HTMRs. In line with our prior work, we identified m1 intradental HTMRs based on their stereotyped fluorescent responses to electrical stimulation (Fig. 1A-B)(*30*, *33*). As expected, intradental HTMRs were robustly activated in response to superficial enamel cutting restricted to the superficial occlusal damage.

**Fig. 1.**
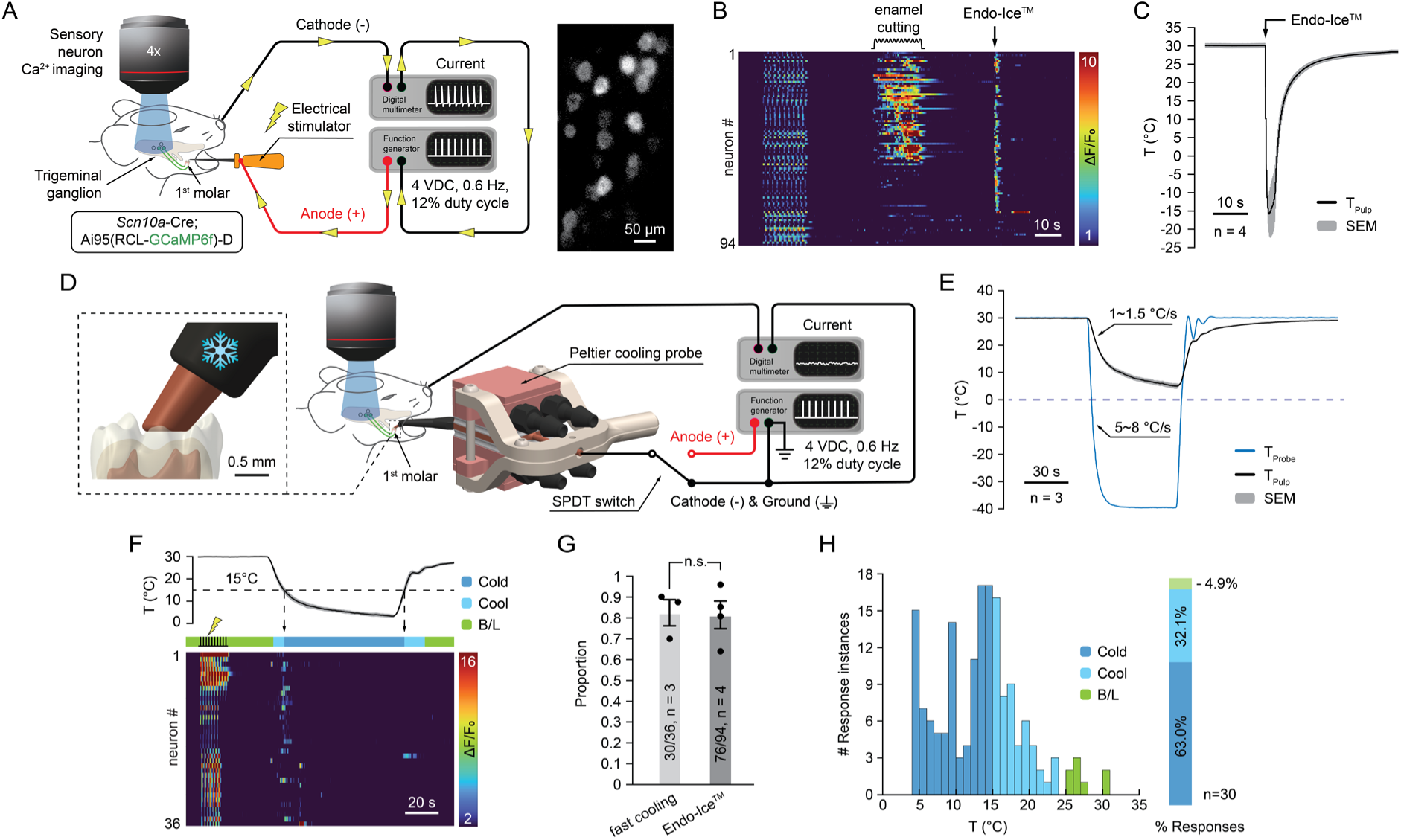
Intradental HTMRs encode cold applied to the molar tooth. **(A)** Schematic of procedure to identify intradental HTMRs during TG in vivo calcium imaging. Application of low voltage pulses to the molar tooth evokes synchronized firing in intradental HTMRs, enabling their rapid identification. Representative epifluorescent image from a *Scn10a*-Cre;GCaMP6f mouse demonstrates the visualization of intradental HTMRs responding to molar electrical stimulation. Scale bar, 50μm. **(B)** Heatmap showing in vivo GCaMP responses from intradental HTMRs tested with select stimuli to the mandibular molar (m1). Intradental HTMRs respond to enamel cutting and Endo-Ice™ application. Data are obtained from *Scn10a*-Cre;GCaMP6f mice. n = 94 intradental HTMRs across 4 TGs. Scale bar, 10s. **(C)** Graph showing measured tooth pulp temperature in an anesthetized mouse during Endo-Ice™ application to the mandibular molar (m1). Measurements are obtained via a thermocouple implanted in the pulp chamber while leaving the occlusal surface of the molar intact. Trace shows the average; shading indicates SEM. n = 4 TGs. Scale bar, 10s. **(D)** Schematic demonstrating the experimental approach to administer precision cooling to an individual molar. The Peltier probe tip was placed on the molar occlusal surface. A thin layer of thermally and electrically conductive silver epoxy was applied between the probe tip and the molar to enhance thermal and electrical transport (see Methods). Administration of electrical stimulation via the probe enabled identification of intradental HTMRs and confirmed sufficient contact prior to administering cooling. **(E)** Graph showing the probe temperature (blue) and corresponding tooth pulp temperature (black) during fast molar cooling. Traces show the average; shading indicates SEM. n = 3 trials. Scale bar, 30s. **(F)** Heatmap showing in vivo GCaMP responses of intradental HTMRs to fast molar cooling. Color bar indicates pulp temperature ranges, including cold (<15°C), cool (15°C∼25°C), and baseline (>25°C). n = 36 intradental HTMRs across n = 3 TGs. Scale bar, 20s. **(G)** Bar graph comparing the proportion of intradental HTMRs responding to fast molar cooling versus Endo-Ice™. Endo-Ice™ responders are analyzed from primary data used in (B). Plotted individual data points represent the proportion of responding intradental HTMRs per trial. Bar shows the mean value and error bars indicate the SEM. n = 3 TGs for fast molar cooling and n = 4 TGs for Endo-Ice™. p = 0.8048 using unpaired t test with Welch’s correction. **(H)** Histogram showing the response instances of n = 30 intradental HTMRs responding to fast molar cooling. The slice plot shows the percentage of firing instances that occurred at cold, cool, and baseline pulp temperatures.

Next, we performed the Endo-Ice™ assay during functional imaging by saturating a small cotton pellet with Endo-Ice™ and immediately applying it to the m1 occlusal surface. We observed strong, transient activation of intradental HTMRs to Endo-Ice™ (Fig. 1B). While it is known that Endo-Ice™ will rapidly cool the enamel surface temperature to -26°C, we sought to determine how this translates to the inner tooth pulp temperature. To this end, we surgically implanted a miniature thermocouple into the pulp chamber while preserving the structural integrity of the occlusal surface (See Methods) and measured temperature while applying Endo-Ice™ in a live, anesthetized mouse. We observed that Endo-Ice™ application drove the internal tooth pulp chamber to -15°C within 1.1 seconds (Fig. 1C), confirming this stimulus results in noxious pulp cooling well below the physiological range. While this data suggests that intradental HTMRs may have the capacity to encode cold temperatures, accurate thresholds and latencies of responses could not be discerned with this rudimentary approach.

### Approach to administer precision cooling to an individual molar

Having demonstrated that intradental HTMRs are activated by Endo-Ice™, we next sought to systematically determine how intradental HTMRs encode low temperatures. To this end, we custom-engineered a Peltier probe capable of cooling an individual molar with precise temperature set points and cooling rates while we conducted our in vivo calcium imaging assays (Fig. 1D). This system was designed with two interchangeable modes that can deliver either electrical or thermal stimuli. To selectively identify intradental HTMRs prior to cooling, we electrically stimulated m1 via the probe tip, thereby enabling functional identification of intradental HTMRs and ensuring sufficient contact between the probe and the occlusal tooth surface. To prevent any potential for static discharge during cooling, we subsequently switched to a grounded configuration prior to thermal stimulation (Fig. 1D). Empirically, we determined the Peltier probe reliably cooled the internal tooth pulp to 2°C at defined cooling rates of 0.1°C/s to 1.5°C/s (Fig. 1E, Fig. S1A-B). Given that intradental HTMR responses observed during Endo-Ice™ application are elicited by rapid cooling to a noxious range (Fig. 1C), we first sought to determine whether they respond to controlled rapid cooling within physiologically relevant low temperatures. Probe-driven rapid pulp cooling (30°C to 5°C at 1∼1.5°C/s) elicited robust, reversible responses from intradental HTMRs (Fig. 1F). These responses were temporary despite the application of persistent cold (5°C) after the downward temperature ramp. Importantly, we observed comparable proportions of intradental HTMRs responding to probe cooling (30/36, 83.3%) vs. Endo-Ice™ (76/94, 80.9%) (Fig. 1G), indicating these methods were similarly efficient in recruiting neuronal activation. Analysis of the number of intradental HTMRs response instances during probe fast cooling revealed that intradental HTMRs fired primarily in the noxious cold range (<15°C, 104/165, 63.0%) (Fig. 1H). Taken together, precision cooling reveals that intradental HTMRs respond to low temperatures with the majority of responses occurring in the noxious cold range, establishing them as polymodal nociceptors capable of encoding both cold and mechanical damage.

### Intradental HTMRs are activated by absolute noxious cold temperatures independent of relative temperature change and cooling rate

Having demonstrated that intradental HTMRs respond to cold stimulation of the tooth, we next sought to capitalize on the spatiotemporal cooling capability of our device to investigate if responses could be tied to independent qualities of cooling. To this end, we designed a series of cooling paradigms to evaluate whether intradental cold responses were triggered by a specific low temperature threshold and/or the rate of cooling.

We began by devising a stepwise-fast cooling paradigm, which was modified from rapid cooling by inserting a temperature plateau at ∼17°C (Fig. S1B). This 40-second plateau allowed the tooth to regain thermal equilibrium at ∼17°C, separating the rapid cooling paradigm into two independent segments with similar temperature step amplitudes: one in the innocuous temperature range (baseline to ∼17°C) and one in the noxious range (∼16°C to ∼1°C). Stepwise-fast cooling broadly elicited intradental cold responses (Fig. 2A), with recruitment across the intradental HTMRs population (23/31, 74.2%) similar to rapid pulp cooling (30/36, 83.3%) (Fig. 2B). Importantly, the majority of intradental HTMR response instances (62/72, 86.1%) occurred during the noxious cold step (Fig. 2C), indicating their tuning to absolute cold temperatures, but not relative temperature change. Interestingly, time constant analysis revealed that the innocuous cool step (*τ* ≈ 3.9s) was ∼2.5x kinematically faster than the noxious cold step (*τ* ≈ 9.8s), further suggesting that a higher cooling rate does not contribute to triggering intradental cold responses (Fig. 2D). Moreover, intradental HTMR responses converged more closely to a threshold temperature during stepwise-fast cooling than during rapid pulp cooling, implying that accounting for intradental thermal equilibrium is crucial for studying the underlying responding temperature. (Fig. 2E)

**Fig. 2.**
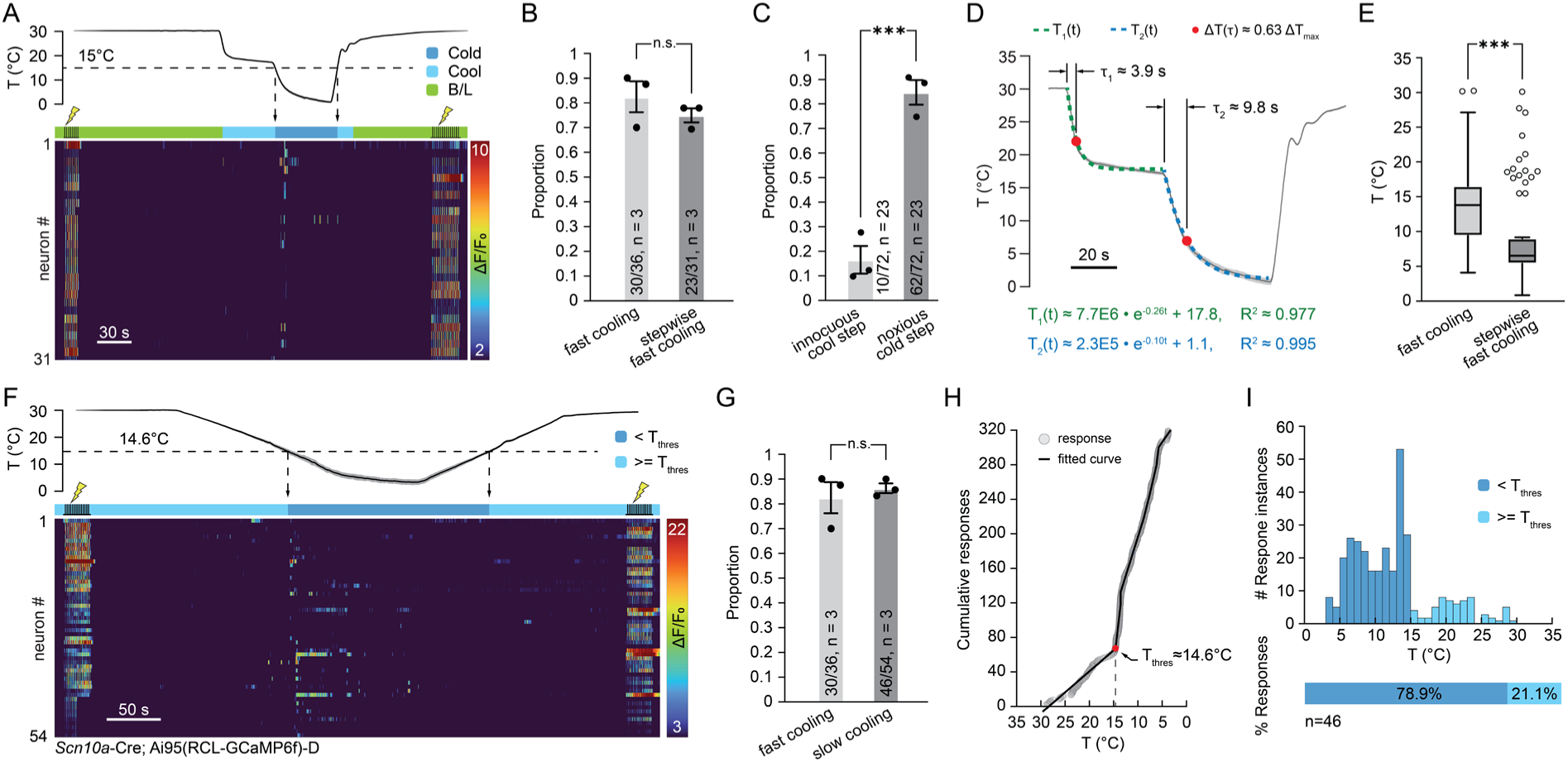
Intradental HTMRs respond to noxious cold independent of relative temperature change and cooling rate. **(A)** Heatmap showing in vivo GCaMP responses of intradental HTMRs to stepwise fast cooling. n = 31 intradental HTMRs across n = 3 TGs. Scale bar, 30s. **(B)** Bar graph comparing the proportions of intradental HTMRs responding to fast cooling versus stepwise fast cooling. Proportions of fast cooling responders are taken from Fig. 1G. Plotted individual data points represent the proportion of responding intradental HTMRs per trial. Bar shows the mean value and error bars indicate the SEM. n = 3 TGs per condition. p = 0.3586 using unpaired t test with Welch’s correction. **(C)** Bar graph comparing the proportions of response instances occurring during the innocuous cool step versus during the noxious cold step across n = 23 intradental HTMRs responding to stepwise fast cooling. Plotted individual data points represent the proportion of intradental HTMR response instances within each temperature step per trial. Bar shows the mean value and error bars indicate the SEM. n = 3 TGs. p = 0.0006 using unpaired t test with Welch’s correction. **(D)** Graph showing thermal kinematics of the two temperature steps in stepwise fast cooling. Trace shows the average pulp temperature; dotted curves represent exponential fit for each step (innocuous cool step: R^2^ ≈ 0.977, noxious cold step: R^2^ ≈ 0.995). Time constant *τ* for each exponential fit represents the time required for the temperature decrease to equal (1-1/e) of its final value. Scale bar, 20s. **(E)** Box plot comparing intradental HTMR response temperatures between fast cooling and stepwise fast cooling. Fast cooling and stepwise fast cooling data are correspondingly analyzed from responding neurons in Fig. 1F and Fig. 2A. Boxes show median and upper/lower quartiles, whiskers indicate maximum and minimum, and circles represent outliers (Tukey). n = 3 TGs per condition. p < 0.0001 using unpaired t test with Welch’s correction. **(F)** Heatmap showing in vivo GCaMP responses of intradental HTMRs to slow cooling. n = 54 intradental HTMRs across n = 3 TGs. Scale bar, 50s. **(G)** Bar graph comparing the proportions of intradental HTMRs responding to fast cooling versus slow cooling. Proportions of fast cooling responders are taken from Fig. 1G. Plotted individual data points represent the proportion of responding intradental HTMRs per trial. Bar shows the mean value and error bars indicate the SEM. n = 3 TGs per condition. p = 0.6096 using unpaired t test with Welch’s correction. **(H)** Graph showing the intradental HTMR threshold response temperature in *Scn10a*-Cre mice. Circles represent the cumulative response instances of intradental HTMRs responding to discrete low temperatures during slow cooling, where slope indicates the frequency of neuronal activations. Line represents a piecewise linear curve fit (up to 10 segments), along which the singularity with the highest slope transition determines the threshold response temperature (14.6°C). n = 54 intradental HTMRs across n = 3 TGs. **(I)** Histogram showing the response instances of n = 46 intradental HTMRs that responded to slow cooling. Slice plot shows the percentage of firing instances that occurred above and below threshold response temperatures calculated in (H).

With evidence suggesting that quasi-static cooling may further resolve the intradental HTMRs’ cold response threshold, we employed a slow cooling paradigm with a pulp cooling rate of 0.1°C/s to determine a putative threshold temperature. Slow cooling effectively triggered intradental cold responses (Fig. 2F) and yielded similar neuronal responsiveness (46/54, 85.2%) to fast cooling (30/36, 83.3%) (Fig. 2G). Noticeably, a substantial increase in neuronal activity (∼1.5 responses/°C to ∼15 responses/°C) occurred at a noxious cold temperature (threshold response temperature, ≈14.6°C) (Fig. 2H). Population response density peaked around this threshold response temperature with the majority (78.9%) of responses occurring below this temperature (Fig. 2I). Taken together, we establish that intradental HTMRs are tuned to encode noxious cold temperatures at an absolute threshold independent of cooling rate. This combined application of precision cooling with intradental functional imaging also establishes, for the first time, detailed insights into how a discrete population of peripheral somatosensory neurons within an internal tissue is tuned to specific features of cooling in vivo.

### Cold responses of intradental HTMRs are independent of Trpm8 or Scn10a (Nav1.8)

Having demonstrated that intradental HTMRs encode noxious cold, we next sought to examine whether these responses rely on their expression of candidate cool or cold molecular mediators. Although some existing evidence indicates that HTMRs comprise a molecularly distinct population from *Trpm8*+ somatosensory neurons(*13*, *16–18*), it cannot be ruled out that *Trpm8* could be expressed at low levels in discrete subpopulations that escape detection in large-scale sequencing(*46*). Furthermore, mixed evidence exists regarding whether intradental neurons express *Trpm8*(*19*, *35*, *47*). To resolve whether intradental HTMRs express *Trpm8*, we performed retrograde tracing of intradental HTMRs using our previously established methods(*30*, *35*) with subsequent multi-plexed ISH. Staining for expected HTMR markers *S100b+* and *Scn10a+* confirmed co-expression in the majority of labeled intradental HTMRs (*S100b+:* 147/147, 100%, *Scn10a+*: 120/147, 82%). Staining for *Trpm8* as well as additional heat-sensing thermoTRP channels *Trpa1* and *Trpv1* revealed no detectable co-expression of labeled neurons with any of these thermoTRP channels (0/147 cells for all 3 markers, 0%). Taken together, our data indicate that noxious cold encoding is not mediated by the canonical cool-sensing receptor *Trpm8* in these neurons. It is worth noting that our finding that intradental HTMRs also lack *Trpv1* expression aligns with a recent report observing minimal expression of *Trpv1* in dorsal root ganglia (DRG) Aδ-HTMRs, confirming similar expression patterns across the two sensory ganglia(*37*). These data indicate that thermosensory encoding by HTMRs throughout the periphery is likely to rely on noncanonical thermosensory mechanisms independent of *Trpm8*, *Trpv1*, and *Trpa1* (Fig. 3A).

**Fig. 3.**
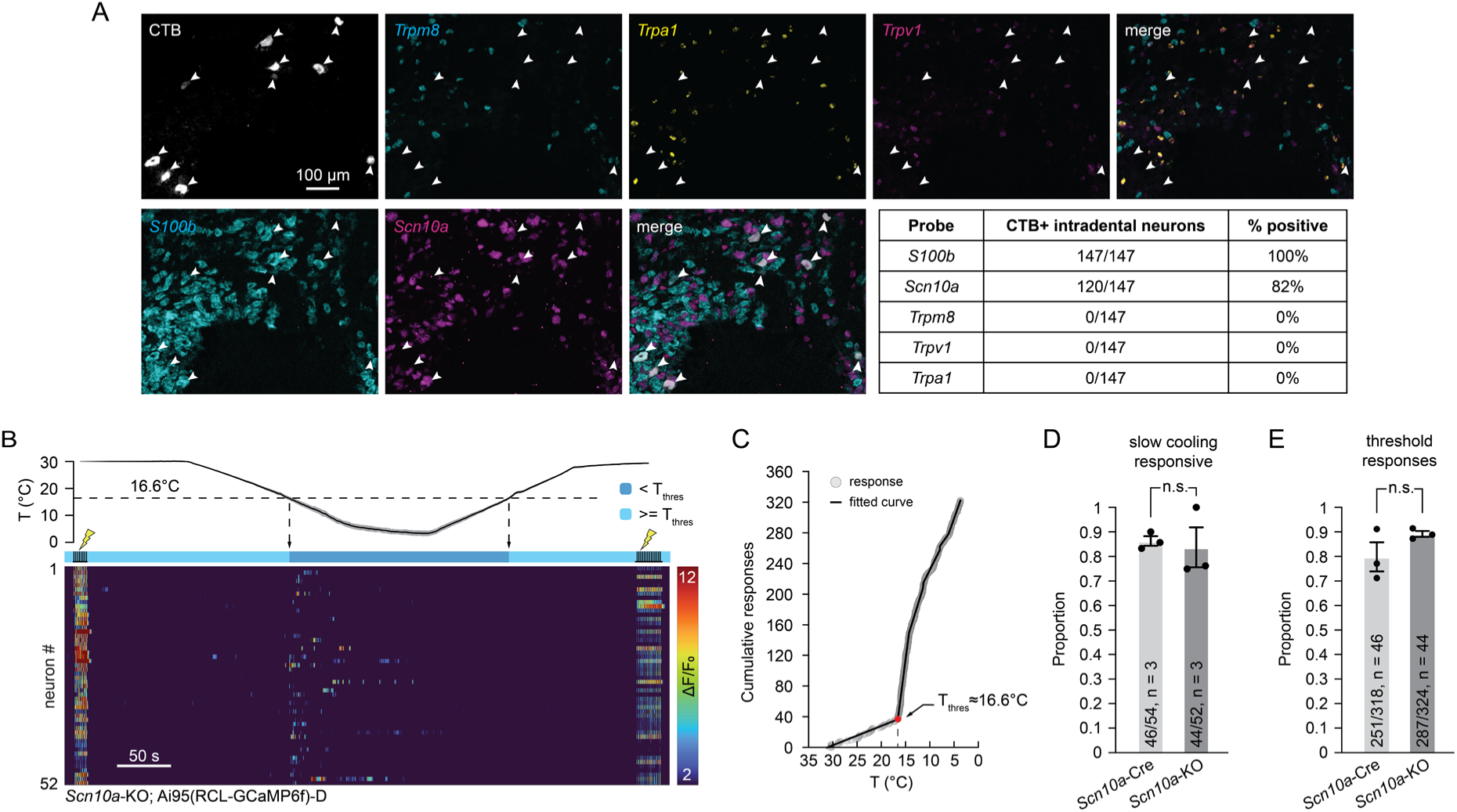
Intradental HTMRs are not canonical cold thermoreceptors. **(A)** Intradental HTMRs do not express thermoTRP channels TRPM8, TRPV1, or TRPA1. Representative image of retrograde-labeled intradental HTMRs (CTB-AF647) and ISH staining. Probes: s100 calcium-binding protein B (*S100b*, cyan), α-Nav1.8 (*Scn10a*, magenta), TRPM8 (*Trpm8*, cyan), TRPA1 (*Trpa1*, yellow), TRPV1 (*Trpv1*, magenta). n = 147 intradental HTMRs. **(B)** Heatmap showing in vivo GCaMP responses of intradental HTMRs to slow cooling in *Scn10a*-KO mice. n = 52 intradental HTMRs across n = 3 TGs. Scale bar, 50s. **(C)** Graph showing the intradental HTMR threshold response temperature in *Scn10a*-KO mice. Circles represent the cumulative response instances of intradental HTMRs responding to discrete low temperatures during slow cooling, where slope indicates the frequency of neuronal activations. Line represents piecewise linear curve fit (up to 10 segments), along which the singularity with the highest slope transition determines the threshold response temperature (16.6°C). n = 52 intradental HTMRs across n = 3 TGs. **(D)** Bar graph comparing the proportions of intradental HTMRs responding to slow cooling between *Scn10a*-Cre and *Scn10a*-KO. Proportions of *Scn10a*-Cre slow cooling responders are taken from Fig. 2G. Plotted individual data points represent the proportion of responding intradental HTMRs per trial. Bar shows the mean value and error bars indicate the SEM. n = 3 TGs per condition. p = 0.7813 using unpaired t test with Welch’s correction. **(E)** Bar graph comparing the proportion of response instances that occurred below threshold response temperatures for *Scn10a*-Cre versus *Scn10a*-KO across n = 46 (*Scn10a*-Cre) and n = 44 (*Scn10a*-KO) intradental HTMRs responding to slow cooling. *Scn10a*-Cre slow cooling data are analyzed from primary data used in Fig. 2F-I. Plotted individual data points represent the proportion of intradental HTMR response instances occurring below the threshold response temperature per trial. Bar shows the mean value and error bars indicate the SEM. n = 3 TGs per condition. p = 0.2521 using unpaired t test with Welch’s correction.

Intradental HTMRs are marked by expression of *Scn10a*, which encodes the voltage-gated sodium channel Nav1.8 that has been implicated in mechano-nociceptive transduction(*48*). A few lines of evidence have implicated a role for Nav1.8+ somatosensory neurons in contributing to cold pain nociception, although conflicting reports suggest this population may not depend on Nav1.8 function(*11*, *12*, *49*, *50*). Here, we investigated whether *Scn10a* is required for intradental HTMR cold responses. We again took advantage of our *Scn10a*-Cre driver line, which is engineered as a knock-in/knock-out driver strain, enabling genetic deletion of *Scn10a* in homozygous alleles while retaining Cre expression(*45*). In *Scn10a*-KO mice, we observed no significant changes in cold responses (Fig. 3B-E), indicating that *Scn10a* and its gene product Nav1.8 are dispensable for noxious cold responses in intradental HTMRs. In summary, we demonstrate that intradental HTMR noxious cold response does not rely on either TRPM8 or Nav1.8. Additional studies are needed to determine whether intradental HTMR cold encoding relies on the presence of additional noncanonical cold receptors.

### Intradental HTMRs are marked by Bmpr1b-Cre

As we determined that intradental HTMRs respond to noxious cold, we next investigated whether this represents a broader capability for this neuronal class to encode cold in other peripheral targets. As our prior molecular characterization of intradental HTMRs demonstrated that they are a homogeneous population of HTMRs marked by high expression of the putative nociceptive marker *Scn10a*(*30*), we were able to employ the *Scn10a*-Cre driver line, despite this line also marking several additional unmyelinated c-type somatosensory populations outside the teeth. Thus, we sought to first identify a more specific Aδ-HTMR Cre line marking intradental HTMRs for functional examination in other peripheral tissues.

Single cell RNA sequencing of DRG somatosensory populations has shown that Aδ-HTMRs can be further subdivided by discrete expression and genetic capture using either *Bmpr1b* or *Smr2*(*13*). It is worth noting that DRG functional imaging using these two lines demonstrated that, at least within cutaneous skin, both are primarily tuned to encode harsh mechanical and heat stimuli(*13*, *37*), although limited data hint that at least a subset of *Bmpr1b*-Cre+ neurons may also encode low temperatures(*13*). Regardless, differences in functional tuning may also exist between the TG and DRG. Therefore, we first set out to determine whether either of these lines might capture intradental HTMRs. To this end, we crossed each line into Ai95D and performed TG calcium imaging to examine whether molar electrical stimulation elicits functional responses. Interestingly, we found that electrical stimulation of the molar tooth drove entrained responses in neurons marked by *Bmpr1b-*Cre (Fig. 4A), but rarely in *Smr2*-Cre (Fig. S2A). As expected, intradental HTMRs found using *Bmpr1b-*Cre also respond to enamel cutting and Endo-Ice™ cooling, indicating that this line faithfully labels intradental HTMRs (Fig. 4A-B). Additionally, we performed histological characterization of intradental terminal endings in *Bmpr1b-*Cre and confirmed robust labeling of terminal endings extending throughout the tooth pulp and inner third of the dentin tubules (Fig. 4C). This labeling and terminal morphology are consistent with our prior histological characterization of intradental HTMRs using *Scn10a*-Cre and *S100b*-Cre driver lines(*30*).

**Fig. 4.**
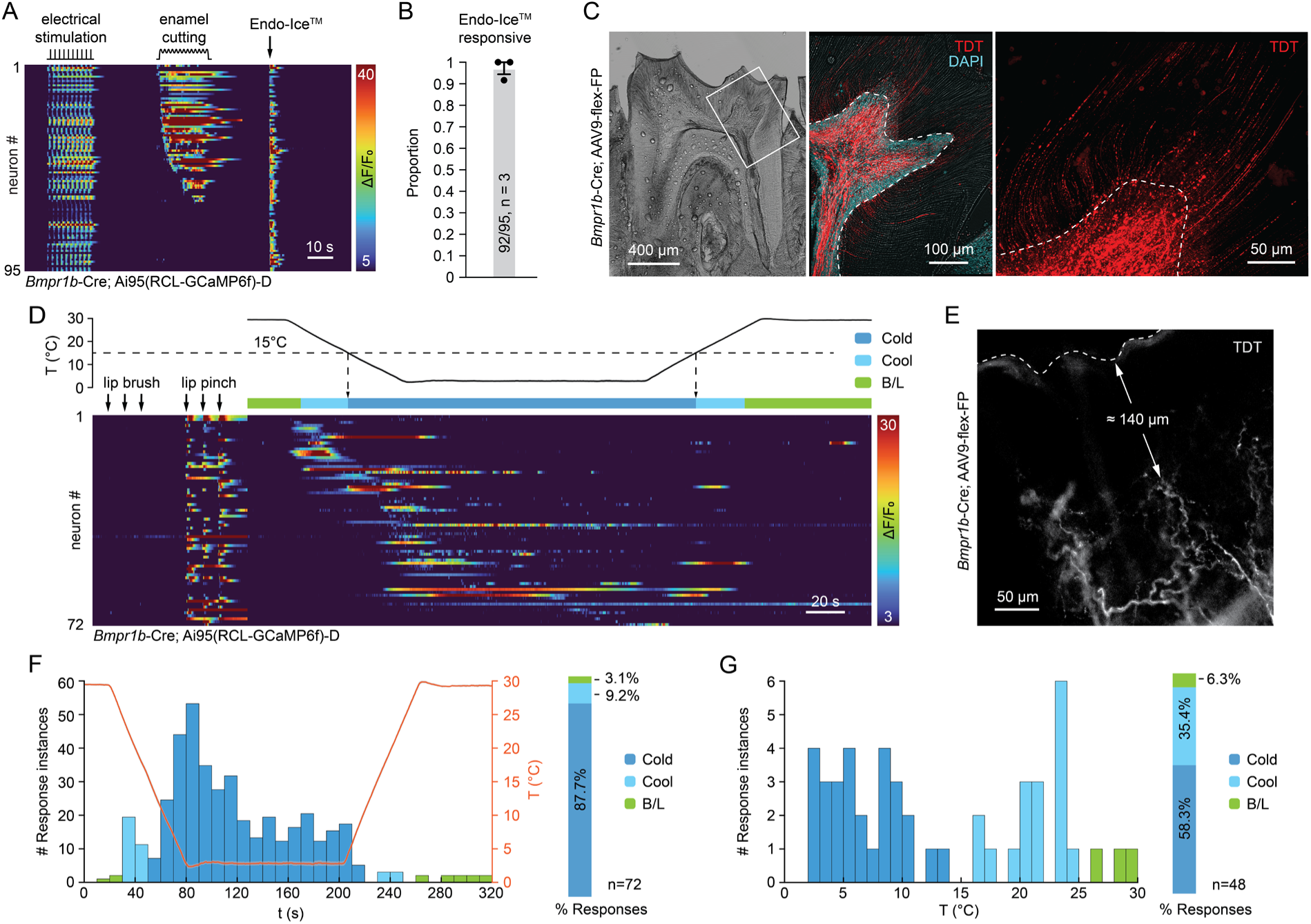
*Bmpr1b*-Cre reveals intradental HTMRs represent a subset of polymodal HTMRs. **(A)** Heatmap showing in vivo GCaMP responses from intradental HTMRs tested with select stimuli to the m1 mandibular molar. Intradental HTMRs respond to enamel cutting and Endo-Ice™ application. Data are obtained from *Bmpr1b*-Cre;GCaMP6f mice. n = 95 intradental HTMRs across 3 TGs. Scale bar, 10s. **(B)** Bar graph showing the proportion of intradental HTMRs responding to Endo-Ice™ in *Bmpr1b*-Cre mice. Plotted individual data points represent the proportion of responding intradental HTMRs per trial. Bar shows the mean value and error bars indicate the SEM. n = 3 TGs. **(C)** Representative image of molar neuronal innervations from *Bmpr1b*-Cre mice receiving intraperitoneal AAV injection of AAV9-Cre. Left panel: Bright-field image of molar cross section. Middle panel: Magnified image of the white rectangle. Merged image overlaid with DAPI staining labeling nuclei of pulp cells, endogenous fluorescence for RFP, and bright-field image. Dashed lines indicate the pulp-dentin border. Right panel: Endogenous fluorescence for RFP is localized to *Bmpr1b*-positive terminal endings that radiate from the pulp and extend into the dentin. This experiment was repeated in n = 4 mice. Scale bars: 400µm (left panel), 100µm (middle panel), or 50µm (right panel). **(D)** Heatmap showing in vivo GCaMP responses of TG neurons responding to lip pinch and slow lip cooling but not to lip brush in *Bmpr1b*-Cre mice. n = 72 neurons across 6 TGs. Scale bar, 20s. **(E)** Representative image of lip neuronal innervations from *Bmpr1b*-Cre mice receiving intraperitoneal AAV injection of AAV9-Cre. Dashed lines indicate the lip surface. Endogenous fluorescence for RFP is localized to *Bmpr1b*-positive terminal endings that extend to the lip surface. This experiment was repeated in n = 3 mice. Scale bar, 50µm. **(F)** Histogram showing TG neurons’ firing instances during slow lip cooling, analyzed from primary data used in (D). Orange trace shows the average lip surface temperature; shading indicates SEM. Slice plot shows the percentage of firing instances that occurred at cold, cool, and baseline lip surface temperatures. n = 6 TGs. **(G)** Histogram showing TG neurons’ first firing instances that occurred before the lip temperature plateau, analyzed from primary data used in (D). Slice plot shows the percentage of first firing instances that occurred at cold, cool, and baseline lip surface temperatures. n = 6 TGs.

### Bmpr1b-Cre HTMRs encode noxious cold in lip soft tissue

We next elected to examine whether trigeminal *Bmpr1b*-Cre-expressing HTMRs mark sensory neuron populations that encode noxious mechanical and cold stimuli in other peripheral tissues. To identify additional peripheral targets receiving innervation from *Bmpr1b-*Cre-labeled HTMRs, we next performed in vivo TG calcium imaging in *Bmpr1b*-Cre;Ai95D and screened for responses to directed tissue pinch, which is known to strongly activate HTMRs. We observed robust neural responses when pinching the ipsilateral mandibular lip (Fig. 4D, S2B). Therefore, we elected to examine whether precision cooling applied to the same ‘pinch’ site also evoked responses. We first set the Peltier probe to the baseline superficial tissue temperature (30°C), brought it into direct contact with the lip, and waited (60s) for thermal equilibration before administering cooling. We reasoned this deliberate delay as well as our validation that HTMRs demonstrate minimal response to gentle brushing (Fig. 4D) would reduce confounding activity. We chose to apply a slow cooling paradigm (0.5°C/s, Fig. S1C) from 30°C to 5°C and examined responses to cooling in neurons that also responded to lip pinch. Intriguingly, we observed 20.3% of pinch-identified neurons also responded during cooling (Fig. 4D, Fig. S2B-C), with most response instances occurring in the noxious cold temperature range (Fig. 4F) and the majority of initial responses arising at noxious cold temperatures (Fig. 4G). We further confirmed that *Bmpr1b-*Cre innervates the lip by performing histology and observed labeled terminal endings in the lip dermis (Fig. 4E). While sequencing data predicts that *Bmpr1b*-Cre is distinct from the *Trpm8*+ neurons(*13*, *16–18*), we further confirmed this via ISH examining for co-expression of *Trpm8* and observed minimal overlap with the genetically encoded reporter (93/1059, 8.8%) (Fig. S2D-E). This indicates that cold responses in these HTMRs are similarly independent of TRPM8. Taken together, these studies show that intradental HTMRs represent a broad subclass of polymodal nociceptors marked by *Bmpr1b*-Cre that are capable of encoding noxious cold and damaging forces in internal and external orofacial tissues.

### Pulp inflammation induces intradental HTMR hypersensitivity

Given that dental pulp inflammation (i.e., pulpitis) causes tooth cold and mechanical hypersensitivity in human patients(*40*), we next sought to determine whether intradental HTMRs activity might correlate with this pain. Importantly, our previous work established that chemogenetic activation of intradental HTMRs drives pain behavior in awake, freely behaving mice(*30*). Thus, cellular activation of intradental HTMRs observed during calcium imaging serves as a proxy for intradental-driven pain. We sought to establish a murine model of early onset pulpitis to investigate whether inflammation alters intradental HTMR responses using in vivo TG imaging. Acute pulpitis was induced in *Scn10a*-Cre; Ai95D mice via application of an inflammatory mediator cocktail (“inflammatory soup”, IS) comprised of well-established acute pain-inducing chemicals(*51*, *52*) to the surgically exposed dentin of m1 in anesthetized mice. Following IS application, a tiny dental filling was placed, and the mouse was allowed to recover for 2 days prior to in vivo imaging (Fig. 5A). To account for potential effects of the surgical process (e.g., dentin exposure, liquid addition), we included a control group receiving 1X PBS to the exposed dentin. Immunohistochemistry staining for the inflammatory marker CD45 demonstrated localized signal enrichment restricted to the IS group, demonstrating successful induction of pulpitis in IS but not PBS control group (Fig. S3).

**Fig. 5.**
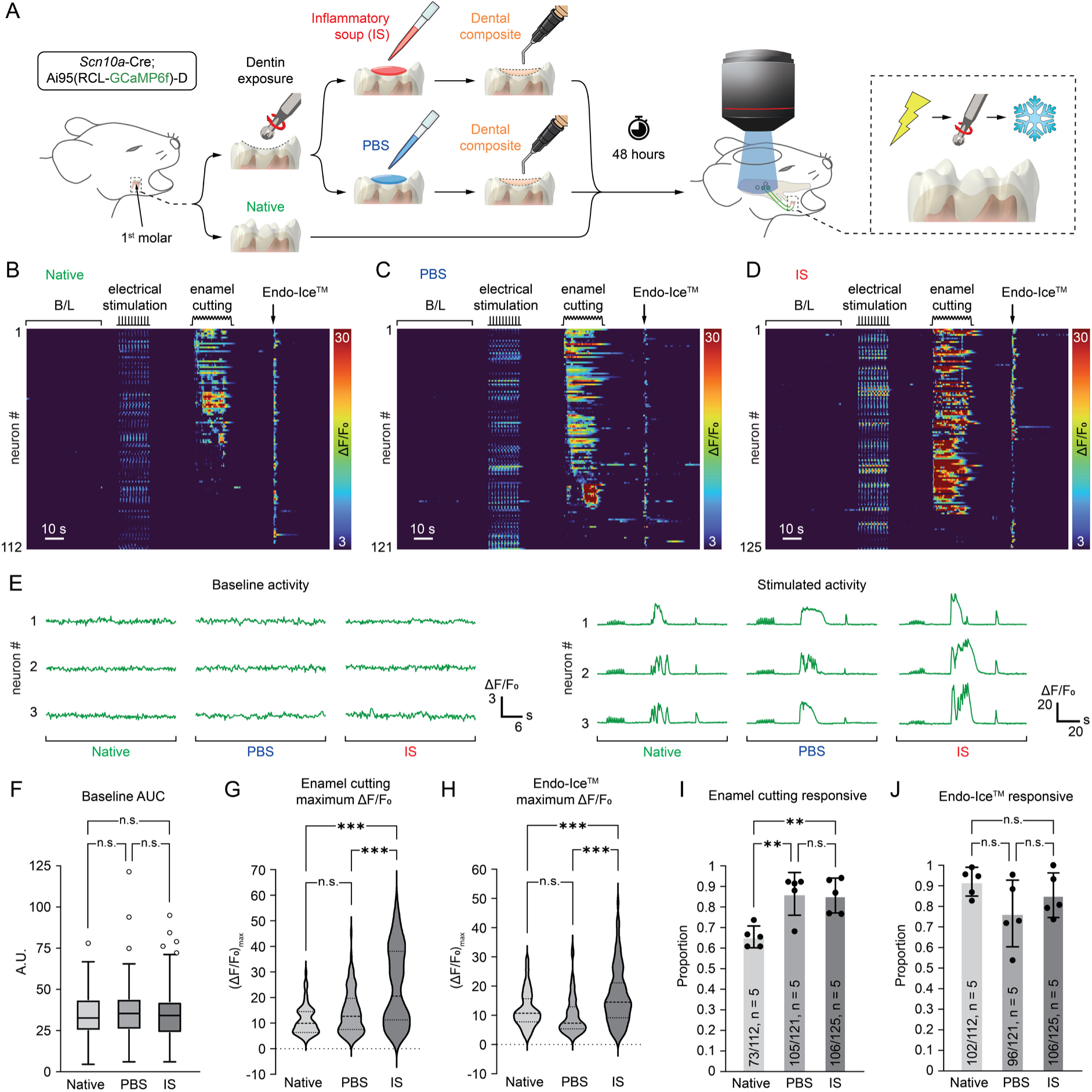
Pulp inflammation sensitizes intradental HTMR response to mechanical and cold stimuli. **(A)** Schematic illustrating the experimental timeline for induction of chronic tooth inflammation of *Scn10a*-Cre;GCaMP6f mice, followed by in vivo TG imaging 48 hours later. Chronic tooth inflammation was induced via preparation of shallow surgical cavitations to expose the mandibular molar (m1) dentin, followed by application of inflammatory soup (IS), which was then sealed with dental composite. Littermate controls either underwent this procedure but instead received PBS, or underwent no surgical procedure (native tooth). **(B)** Heatmap showing the effect of IS-induced chronic inflammation on intradental HTMR responses at baseline and in response to enamel cutting and Endo-Ice™. n = 125 intradental HTMRs across 5 TGs. Scale bar, 10s. **(C)** Heatmap showing intradental HTMR responses at baseline and following control PBS application in littermate controls. n = 121 intradental HTMRs across 5 TGs. Scale bar, 10s. **(D)** Heatmap showing intradental HTMR responses of a native molar at baseline and in response to enamel cutting and Endo-Ice™ in littermate controls. n = 112 intradental HTMRs across 5 TGs. Scale bar, 10s. **(E)** Example of GCaMP traces for individual HTMRs from each experimental condition showing baseline (left) and stimulated (right) activities. Scale bar: 3*ΔF*⁄*F*_0_ (left, vertical), 6s (left, horizontal), 20*ΔF*⁄*F*_0_ (right, vertical), 20s (right, horizontal). **(F)** Box plot showing quantification of unstimulated activity. Boxes show median and upper/lower quartiles, whiskers indicate maximum and minimum, and circles represent outliers (Tukey). n = 5 TGs per condition. p = 0.7375 (IS vs. PBS), p = 0.8152 (IS vs. native tooth), and p = 0.3808 (PBS vs. native tooth) using one-way ANOVA with Tukey’s correction. **(G and H)** Violin plots showing the maximum response intensity of intradental HTMRs responding to (G) enamel cutting and (H) Endo-Ice™ in each condition. Results in (G) and (H) are from primary data used in (B-D). n = 5 TGs per condition. For enamel cutting, p < 0.0001 (IS vs. PBS), p < 0.0001 (IS vs. native tooth), and p = 0.1426 (PBS vs. native tooth) using one-way ANOVA with Tukey’s correction. For Endo-Ice™, p < 0.0001 (IS vs. PBS), p < 0.0001 (IS vs. native tooth), and p = 0.1401 (PBS vs. native tooth) using one-way ANOVA with Tukey’s correction. **(I and J)** Bar graphs showing the proportion of intradental HTMRs responding to (I) enamel cutting and (J) Endo-Ice™ in each condition. Plotted individual data points represent the proportion of responding intradental HTMRs in each condition. Bar shows the mean; error bars indicate the SEM. Results in (I) and (J) are from primary data used in (B-D). n = 5 TGs per condition. For enamel cutting, p = 0.9842 (IS vs. PBS), p = 0.0064 (IS vs. native tooth), and p = 0.0047 (PBS vs. native tooth) using one-way ANOVA with Tukey’s correction. For Endo-Ice™, p = 0.4895 (IS vs. PBS), p = 0.6698 (IS vs. native tooth), and p = 0.1444 (PBS vs. native tooth) using one-way ANOVA with Tukey’s correction.

To investigate whether and how inflammation shapes HTMR responses, we elected to perform our established stimulation paradigm consisting of molar enamel cutting and Endo-Ice™ application while examining intradental baseline activity as well as responses across all 3 experimental groups (native tooth, PBS, and IS). Intradental HTMR spontaneous activity was indistinguishable between pulpitis or PBS controls (Fig. 5B-F). However, evoked stimulation revealed a distinct divergence in responses. Specifically, IS-induced inflammation resulted in a significant increase in intradental HTMR maximal firing intensity to both enamel cutting (Fig. 5G) and Endo-Ice™ application (Fig. 5H) compared to PBS control or native tooth conditions. This indicates that tooth inflammation potentiates evoked intradental responses without changing their spontaneous activity, providing a putative neural functional basis for latent, evoked tooth hypersensitivity in clinical pulpitis.

Interestingly, while the majority of electrically responding intradental HTMRs respond to enamel cutting in native tooth controls (73/112, 65.2%), we observed a significant increase in the proportion of enamel cutting responders in both PBS (105/121, 86.8%) and IS (106/125, 84.8%) groups (Fig. 5I). As the proportion of responders was similarly elevated in both PBS and IS groups, this indicates that enamel structural damage induced by surgical dentin exposure alone increases the proportion of intradental responses to enamel cutting, suggesting increased mechano-sensitivity. Notably, this proportional increase did not emerge in response to Endo-Ice™ application across all 3 conditions (Fig. 5J), suggesting potential mechanistic differences between intradental cold- and mechano-sensation. In summary, our findings reveal a somatosensory basis for the neural origins of pulpitis-induced tooth pain, and demonstrate the functional logic for pulpitis-induced tooth hypersensitivity.

## Discussion

Based on both clinical observation that tooth sensation is closely tied to noxious cold pain and our recent work demonstrating that intradental innervation is predominantly composed of myelinated HTMRs that drive pain, we set out to determine whether this population also encodes cold within the context of the mammalian tooth. To uncover fundamental mechanisms of peripheral cold encoding in this system, we incorporated a powerful combination of dynamically controlled spatiotemporal cooling with in vivo calcium imaging to probe sensory neurons for their ability to encode cold (<15°C) in both the tooth and orofacial soft tissue. We conclude that intradental HTMRs: 1) are the primary neural mediator of noxious cold; 2) respond to an absolute cold temperature threshold independent of the rate of temperature change; 3) do not rely on canonical cool sensor TRPM8 and putative cold pain transducer Nav1.8; 4) are marked by the HTMR subpopulation driver line *Bmpr1b*-Cre but not *Smr2*-Cre; and 5) demonstrate exaggerated mechanical and cold responses, but not de novo spontaneous activity, in the context of pulpitis. Outside the tooth, our work further implicates HTMRs in peripheral craniofacial cold encoding based on calcium imaging of neurons marked by *Bmpr1b*-Cre. Taken together, our work provides significant new insights to understanding this dental phenomenon and the neural basis for cold sensation.

Inner tooth innervation provided a unique opportunity to characterize the role of myelinated HTMRs in cold encoding because it represents a relative enrichment of a homogenous neural class (HTMRs). Our work establishes these HTMRs as primary neural mediators within the teeth that encode noxious cold and damaging forces. Importantly, we confirmed that intradental HTMRs do not express TRPM8, consistent with transcriptomic data(*13*, *16*, *18*, *35*), demonstrating that this organ encodes cold in the tooth via a TRPM8-independent sensory neuron population. Notably, we subsequently show that the somatosensory neurons marked by *Bmpr1b*-Cre which lack *Trpm8* also represent the intradental HTMR population. Our findings here expand our understanding of the *Bmpr1b*-Cre tool and its ability to mark intradental HTMRs (and potentially HTMRs, broadly) through developmental expression and/or low-level expression that escaped detection using in situ hybridization(*30*). It is worth noting that only a portion of *Bmpr1b*-Cre labeled HTMRs responding to lip pinch also responded to noxious cold (∼20.3%, Figure S2B-C), suggesting the presence of additional molecular heterogeneity. Importantly, in both tooth and lip, HTMRs respond to cold at a threshold much lower than reported for TRPM8(*6*), supporting a mechanism independent of *Trpm8* and suggesting neural populations independent of *Trpm8*+ neurons mediate somatosensory noxious cold encoding. Indeed, our data suggest that this newly identified HTMR population may serve as a signaling partner to second-order Sp5 brainstem neurons in TRPM8-knockout mice during oral cavity cooling(*21*).

By incorporating customized Peltier devices to deliver precision spatiotemporal cooling to individual orofacial tissues during TG functional imaging, we systematically controlled temperature ramp rates and setpoints, revealing that intradental HTMRs encode noxious cold at an absolute temperature threshold independent of relative temperature drops or cooling rates. While human psychophysical studies demonstrate that both absolute thresholds and temporal dynamics can shape thermal perception(*53*, *54*), our data suggest trigeminal myelinated HTMRs contribute to perceptual detection of absolute noxious cold temperatures once a threshold is reached, regardless of temperature dynamics. Our data demonstrate that trigeminal myelinated HTMRs elicit acute responses while cooling below their threshold response temperature and rapidly desensitize despite application of persistent cold temperature. This is in alignment with recent studies of higher-order neurons in the spinal cord(*22*), thalamus, and cortex, which suggest low-temperature encoding is tuned primarily to the temperature-changing phase occurring below the neurons’ activation thresholds. Together, our findings support a mechanism by which this HTMR population drives pain upon their activation at a specific absolute threshold, providing the functional basis for their nociceptive role.

Our work has implications for longstanding models of tooth sensation. In addition to direct sensing of neuron terminal endings to sensory stimuli, two theories founded in the tooth’s unique structural and cellular composition have been proposed to contribute to tooth thermal sensation(*55*). Intradental HTMRs innervate the living tooth pulp and penetrate into overlying mineralized, fluid-filled dentin tubules(*55*, *56*). This tissue organization has inspired the conception of hydrodynamic theory, which posits that tooth sensation arises not directly from discrete sensory stimuli but from the movement of dentinal fluid induced by those stimuli(*55*, *57–60*). Such fluid movement, in turn, generates wall shear stress coupled with fluid velocity, which could mechanically activate primary sensory cells(*61*, *62*). In this model, which preceded the identification of molecular thermoreceptors, dentinal fluid movement is hypothetically driven by temperature-change-induced thermal expansion in both the tubule wall and the dentinal fluid, with increasing rate of temperature change accelerating dentinal fluid movement and resulting in higher wall shear stress(*61*, *62*). In all, according to hydrodynamic theory, higher rates of temperature change are more likely to trigger and exaggerate intradental cold responses. However, hydrodynamic theory fails to explain the results of our spatiotemporal cooling experiments. For one, in the stepwise-fast cooling paradigm, most intradental cold responses occur during the noxious cooling step rather than the innocuous step, despite the latter having faster thermal kinetics to drive fluid movement at higher velocity. Moreover, the slow cooling paradigm elicits robust intradental cold responses despite using a cooling rate significantly slower than either phase of stepwise-fast cooling. Finally, the transient onset of intradental cold responses at intermediate points during any ramps does not correspond to any dynamics of dentinal fluid movement. Together, our study suggests a functional decoupling between fluid movement and sensory tuning in intradental cold sensation.

A second intradental sensation model suggests odontoblasts, which are columnar cells that line the pulp border and function to secrete and maintain mineralized dentin, could function as primary sensory cells that function upstream of the terminal endings to directly transduce stimuli and signal to intradental HTMRs(*55*). This theory is largely based on in vitro evidence due to technical challenges in directly assessing odontoblast excitability in vivo(*63–68*). While our study cannot rule out potential contributions of odontoblasts in the tooth sensory apparatus, our finding that *Bmpr1b*+ HTMRs functionally respond to low temperatures within both the teeth and lip peripheral soft tissue, which are structurally and cellularly distinct tissues, supports that the neuronal terminals may be capable of directly encoding noxious cold in the absence of accessory cells.

Our study also underscores the importance of examining thermosensory tuning in vivo. While in vitro studies are useful and critical for characterizing parameters mediating specific activation of molecular receptors to thermal stimuli(*6*, *8*, *9*), these experiments are performed in an isolated environment that cannot fully recapitulate how cellular mechanisms function in vivo. This is also exemplified by the inability to recapitulate the role of TRPM8 and TRPA1 in noxious cold sensing in vivo at the behavioral level(*9*, *10*, *69–71*) despite prior in vitro characterization of their channel gating(*6*).

Despite the pervasiveness of patients presenting with tooth inflammatory hypersensitivity within dental clinics, the underlying neural mechanisms have remained largely unknown. Here, we determine that intradental HTMRs function as primary neural drivers for this phenomenon within the teeth. We observed heightened evoked responses during acute inflammation that align with recent findings in additional trigeminal peripheral target tissues including cutaneous cheek skin(*15*) and the masseter muscle(*51*), hinting at a conserved functional basis for inflammatory-mediated orofacial pain. Notably, acute cheek skin inflammation also resulted in significant increases in baseline unstimulated activity in both c-type nociceptors and A*δδ*-HTMRs(*15*); in contrast we did not observe spontaneous baseline firing in A*δδ*-HTMRs in pulpitis, suggesting tissue-specific differences. Our finding that the intradental HTMRs drive pulpal sensitivity also explains the paradoxical period where patients lack evoked sensitivity when the pulp is necrotic, but extradental tissues are not yet affected. Thus, our data serve as mechanistic insight explaining these (unfortunately) common clinical phenomena. While we reported heightened activation to both thermal and mechanical stimuli under pulpitis, future work is needed to determine whether the threshold of tuning to these stimuli is affected. Together, these insights may enable the development of more objective diagnostic approaches to functionally distinguish reversible versus irreversible pulpitis.

In summary, our results reveal a new role for trigeminal myelinated HTMRs in encoding absolute noxious cold temperatures in both hard and soft orofacial tissues, and provide the neural and functional basis for inflammatory-mediated hypersensitivity within the teeth, which may represent a common principle across peripheral tissues.

## Acknowledgements

We are grateful to Colleen Mahoney and Charlotte Mistretta for critically reading the manuscript and providing valuable insights, and Dr. David Ginty for *Smr2*-Cre and *Bmpr1b*-Cre knock-in mice.

## Author Contributions

Conceptualization: E.A.R., S.W., K.P.P., and J.J.E.

Methodology: E.A.R., S.W., S.K.P., A.R.G., A.J., K.P.P., and J.J.E.

Investigation: E.A.R, S.W., A.J., S.K.P., A.R.G., J.X., S.T.W., and L.L.W.

Visualization: S.W., E.A.R, A.J., K.P.P., J.J.E.

Funding acquisition: E.A.R., J.J.E.

Project administration: E.A.R., K.P.P., J.J.E., S.W.

Supervision: K.P.P., J.J.E., E.A.R., S.W., S.K.P.

Writing – original draft: E.A.R., S.W., J.X.

Writing – review & editing: E.A.R., S.W., K.P.P., J.J.E.

## Data, code, and materials availability

All original code is deposited at Zenodo and is publicly available at the date of publication. All data and information required to reanalyze the reported data will be shared by the lead contact upon request.

## Supplementary Materials

### Materials and Methods

#### Animal assurance statement

Animal experiments were conducted following approved protocols by the University of Michigan Institutional Animal Care and Use Committee (Protocol Number: (PRO00012299) in alignment with NIH guidelines. Both male and female mice were used for all experiments with sample sizes for each experiment described in the figure legends. Mice were group housed at room temperature on a 12 hour light/ 12 hour dark cycle with *ad libitum* access to water and standard laboratory mouse pellet food. Mouse lines were obtained from Jackson or listed contributors and crossed into reporter lines with no further backcrossing.

#### Mouse lines

The mouse lines used in the present study were: Nav1.8-Cre (#036564, JAX), Ai95(RCL-GCaMP6f)-D (#028865, JAX), C57BL/6J (#000664, JAX), *Smr2*-Cre (#039562), *Bmpr1b*-Cre (#039561).

#### Peltier probes

##### Molar cooling probe

A custom Peltier probe was designed and fabricated to administer temperature and electrical stimuli to an individual mouse molar. The probe assembly adopted a symmetric sandwich structure, with (away from the center plane): a CNC-fabricated copper cooling concentrator, two Peltier modules (25×25×3mm, 04811-5P31-15CK, Custom Thermoelectric) for temperature regulation, and two water blocks (25.4×25.4×12.4mm, WBA-1.00-0.49-CU-01, Custom Thermoelectric) for waste heat dissipation.

The cooling concentrator consists of substructures including a 25x25x2mm core plate, a 2mm OD electrical terminal rod, and a tapered cooling tip with ∼0.4 mm^2^ elliptical tip area. Among these substructures, the core plate contacts the Peltier modules to relay temperature changes, the electrical terminal rod receives external electrical potential control, and the tapered cooling tip transfers thermal and electrical stimuli to an individual mouse molar.

A miniature NTC thermistor (GA10K3MCD1, TE Connectivity) was used to measure probe tip temperature and to enable feedback control via a TEC controller (TCM-M115, Yexian). The thermistor was mounted on the cooling tip using thermal adhesive (K-5205, Kafuter), 2mm offset from the tip point. Insulative varnish (4226A, MG Chemicals) was coated around the thermistor to avoid moisture degradation. The cooling tip and the thermistor were further thermally and electrically insulated with heat-shrink tubing, leaving 1mm of the tip exposed.

To minimize interfacial thermal resistance, thermal sheets (TS-MFK6, Custom Thermoelectric) were inserted between the core plate, Peltier modules, and water blocks. A 3D printed bracket secured the probe assembly to a manipulator, enabling both coarse and fine spatial alignment. During probe operation, a circulating chiller (Haake A40 AC150 1555408, Fisher Scientific) cycled -15°C DD water/antifreeze (Pink, Peak) mixture (60/40) through the probe’s water blocks to facilitate rapid temperature ramping. This custom Peltier probe provided rapid, stable probe-tip temperature control from -40°C to 50°C, with maximum temperature ramp rates of 8°C/s and a steady-state precision of ±0.05°C.

##### Lip cooling probe

The lip cooling probe was modified from the molar cooling probe, which embodied a different cooling concentrator design. The lip cooling probe’s cooling concentrator consists of substructures including a 25x25x3mm core plate, a discharge screw, and a tapered cooling tip with a 2x5mm rectangular tip area. The thermistor (GA10K3MCD1, TE Connectivity) was embedded in a channel 1mm beneath the tip surface using thermal adhesive (K-5205, Kafuter). A ground wire was connected to the cooling concentrator via the discharge screw to eliminate static discharge. This custom Peltier probe provided rapid, stable probe-tip temperature control from 0°C to 50°C, with maximum temperature ramp rates of 10°C/s and a steady-state precision of ±0.05°C.

##### In vivo pulp temperature measurement

In vivo pulp temperature change in response to Endo-Ice^TM^ application or Peltier probe cooling was determined using our previously published procedure to monitor internal pulp temperature with slight modifications(*30*). Briefly, the mouse was anesthetized with isoflurane (4% for induction;1.5∼2% for maintenance) via a SomnoSuite Low-Flow Anesthesia System (Kent Scientific). Immediately after anesthesia induction, ophthalmic ointment (Fisher Scientific Catalog #NC0490117) was applied to both eyes to prevent corneal drying. The mouse was then head-fixed using a custom stereotaxic stage, during which isoflurane was administered via a custom-designed nose cone to maintain a deep plane of anesthesia. Body temperature was maintained using a hand warmer. The mouse’s mouth was opened both vertically and horizontally using a metallic, spring-loaded incisor retractor (anchored on mandibular and maxillary incisors) and a 3D printed buccal retractor. The tongue was gently drawn laterally with a Dieffenbach Serrefine clamp (2.5”) to expose the mandibular molars.

A channel was drilled on the mesial side into a mandibular m1 molar using a needle carbide dental bur (Dentsply) driven by a 35,000RPM micromotor handpiece. The channel extended halfway through m1 to expose the inner pulp chamber without compromising the occlusal tooth structure. A T-Type thermocouple with 50μm wire diameter was then implanted through the channel into the inner pulp chamber, with its bead positioned at the pulp-dentin interface. The channel was then sealed with dental filling (Flow-It ALC B2, Pentron) and UV-cured to secure the thermocouple. Analog thermocouple temperature readings were captured with a thermocouple data logger (TC-2000 Type T Thermocouple Meter, Sable Systems International) with 0.01°C resolution and 0.2°C accuracy. Analog thermocouple data were converted to digital using a DAQ system (LABTRAX 4-Channel Data Acquisition System, WPI Data Acquisition). Thermocouple data were acquired with LabScribe V4 software (IWORX) at 100Hz. The pulp temperature recording protocol involved the full duration of either Endo-Ice™ application or individual Peltier probe cooling paradigms (fast, stepwise fast, slow), with 30-second baseline periods before and after. This protocol ensured reproducible assessment of pulpal thermal dynamics in response to molar cooling.

#### In vivo epifluorescence calcium imaging of the trigeminal ganglion

Trigeminal ganglia (TG) calcium imaging was conducted in vivo using anesthetized mice via published methods with modifications(*30*, *72*). Anesthesia and head-fixation was first achieved following the steps described in the “pulp temperature measurement” section. To eliminate accidental TG neuron activity due to whisker deflection, whiskers were first trimmed close to the mouses face (∼1mm). Surgical exposure of the TG surface was performed via bilateral hemispherectomy followed by achievement of hemostasis, after which the surface of the ganglia could be imaged for ∼30 minutes to 1 hour for optimal neuron responses. GCaMP6f (480nm) imaging was performed using a custom-built Thorlabs epifluorescence microscope equipped with a 4x, 0.16 NA Olympus objective and an Ultra-High-Power LED light source (UHP-F-WDH, Prizmatix), filtered with a GFP filter cube (Thorlabs). To accommodate TG curvature and capture a deeper field of view (300μm) of the ganglion surface, a piezoelectric actuator (NanoScan NPC-D-6111, Prior) was used to provide precise periodic (5Hz) vertical motion of the objective. Timelapse image stacks were acquired with a pco.panda 4.2 bi scientific CMOS camera at a 5Hz acquisition rate.

##### Molar stimuli

Access to the mouse mandibular molars was first obtained using our published method(*30*, *33*). In brief, the mouse’s mouth was held open both vertically and horizontally using a metallic, spring-loaded incisor retractor (anchored on mandibular and maxillary incisors) and a 3D printed buccal retractor. The tongue was gently retracted laterally via a Dieffenbach Serrefine clamp (2.5”) to expose the mandibular molars.

Intradental HTMR somas within the TG were functionally identified via their entrained responses to low DC voltage pulses (“electrical stimulation”) using our previously established protocol(*30*, *33*). For paradigms involving enamel cutting and Endo-Ice™ cooling, an electrical stimulator was first placed on the molar occlusal surface. A single-post double-throw (SPDT) switch was configured to connect the electrical stimulator to the anode position of the electrical stimulation apparatus. The cathode of an electrical stimulation apparatus was connected to the incisor retractor to regulate and monitor electric current exiting the mouse. Current data were timestamped and acquired as references. Enamel cutting was then administered to the occlusal surface of the molar via a ¼ round carbide dental bur (Munce discovery burs, CJM Engineering) driven by a micromotor handpiece (35,000RPM) in a posterior-to-anterior sweeping motion. Endo-Ice™ cooling was applied to the electrically stimulated molar by spraying Hygenic Endo-Ice™ Pulp Vitality Refrigerant Spray (Coltene) onto a cotton ball (1mm diameter) held with forceps for 5 seconds, followed by immediate application.

For paradigms involving Peltier probe molar cooling, a small drop of thermally and electrically conductive silver epoxy (8331D-14G, MG Chemicals) was applied to the probe tip, followed by precise alignment to ensure site-specific contact between the probe tip and the molar occlusal surface. A mild downward force, indicated by minor jaw descent, was applied on the probe tip to spread the silver epoxy. To perform molar electrical stimulation via the Peltier probe tip for intradental HTMR identification and probe-molar contact validation, the SPDT switch was configured to the anode position. The cathode was connected to the incisor retractor to regulate and monitor electric current exiting the mouse. Probe cooling was script-controlled for consistency. During probe cooling, to eliminate potential contamination from static discharge, the SPDT switch was set to the cathode position to equalize the electric potential between the mouse and the probe tip. Post-cooling electrical stimulation was performed by setting the SPDT switch back to the anode position to validate neuron viability. Probe settings, probe temperature, and current readout were timestamped and acquired from the Ca^2+^ imaging computer for synchronization.

##### Lip stimuli

The mouse’s lip was gently stroked with the tip of a fine paintbrush (size 000). Each stroke was cautiously applied to a localized lip region at a consistent force, indicated by slight brush hair deflections. Lip pinch was performed with precision forceps (Dumont #5/45 Forceps, FST) by pinching superficially at the stroked lip region. Lip cooling was administered to the same lip region using a Peltier probe with an enlarged tip contact area (2mm×5mm). A thin layer of thermal paste (Arctic Silver 5) was evenly applied between the probe tip and the lip surface to improve heat transfer. A T-Type thermocouple with 50μm wire diameter as sandwiched between the probe tip and the lip surface to acquire the lip surface temperature. Probe temperature and thermocouple temperature were timestamped and recorded.

#### Calcium imaging analysis

##### Pre-processing of in vivo calcium response time-lapse image data

Image stacks from a single imaging trial were first concatenated in ImageJ (Image - Stacks - Tools - Concatenate). Rigid registration was performed using a previously published MATLAB script to correct for minor inter-frame movements(*30*). For image stacks exhibiting warping, a combination of rigid and nonrigid registration was performed using Suite2p(*73*), a previously published image-processing pipeline. The motion- and warp-corrected image stack was then cropped to remove the uneven edges and photobleach-corrected in ImageJ (Image - Adjust - Bleach Correction) using the exponential fit method.

##### Regions of interest (ROIs) selection

Regions of interest (ROIs) were manually traced with ImageJ’s freehand selection tool. For molar stimulation paradigms, the baseline (average intensity of the first 30 frames prior to stimulation) was subtracted from the electrical stimulation paradigm timelapse to enhance GCaMP6f fluorescence signal contrast. ROIs were drawn around neurons that synchronously responded to the applied electrical stimulation. For lip stimulation paradigms, the raw image stack was split into ‘pre’ and ‘post’ substacks separated by 25 frames (5s). Substack ‘post’ was subtracted from ‘pre’ to enhance fluorescence increase during this 5-second window. ROIs were drawn around neurons that increased in fluorescence in response to the administered mechanical and thermal stimuli.

##### Fluorescence dynamics

Normalized, background-subtracted GCaMP6f fluorescence (*ΔF*⁄*F*_0_) was calculated using a previously published MATLAB script(*14*, *30*), allowing for parallel comparison between neurons with different levels of GCaMP6f expression. For each somatic ROI, background intensity was calculated from the surrounding neuropil region, where contaminant signals from adjacent cells/blood vessels were thresholded out. Background-subtracted somal fluorescence intensity (*ΔF*) was calculated by subtracting background intensity from average intensity inside the ROI. Baseline somal fluorescence intensity (*F*_0_) was calculated as the average of the ten lowest mean somatic intensities. *ΔF*⁄*F*_0_ calculation was performed framewise throughout the image stack.

##### Response instance detection

Response instance detection was performed using MATLAB’s local maxima finding function findpeaks(). For each normalized GCaMP6f fluorescence trace, local maxima outside the stimulation window were first located. From these maxima, mean values and standard deviations of peak heights and prominences were used to reference baseline noise. Response instances were identified based on their peak heights and prominences compared to baseline noise, which followed two criteria:

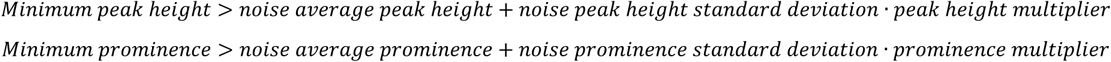

where 3 was used as the prominence multiplier, and either 4 or 6 was adopted as the peak height multiplier to accommodate degree of stimulation differences between paradigms and response intensity differences between mouse lines.

##### Responder determination

Responders were determined in MATLAB using a previously published method(*74*). Neuronal response instances that occurred outside the stimulation window were considered as “off” responses. Neurons with more than one off response were categorized as spontaneous responders and were excluded from statistics.

##### Calculation of proportion of responders

The proportion of determined responders was calculated as the fraction of responders to a specific stimulation paradigm (e.g., Peltier probe cooling, enamel cutting, Endo-Ice^TM^) among all electrical stimulation responders (i.e., intradental HTMRs).

##### Temperature interpolation & alignment

Frame-specific probe temperature was determined using a custom MATLAB script. An equally spaced array of video timestamps was generated from the video’s length (in frames) and the timestamps of the first and last frames. Timestamp and temperature data were extracted from the probe/thermocouple temperature data file and were linearly interpolated using MATLAB’s 1-D data interpolation function interp1() to match the video timestamps.

Frame-specific pulp temperature was then determined using a separate MATLAB script, which aligns the pulp-probe temperature correlation data (∼10Hz) with the interpolated probe temperature data (5Hz). Since probe temperature paradigms were script-controlled and highly consistent, data points between the two furthest probe temperature adjustment points were used as a scaling reference. For each dataset to be aligned, a linearly spaced array with the same start and end values was overlaid on the reference data and extrapolated bidirectionally to span the entire dataset. This array was then used as the scaling input of MATLAB’s interp1() function to map pulp temperature to interpolated probe temperature.

##### Time constant

Time constants for the two stepwise fast cooling segments were calculated using a custom MATLAB script. An exponential curve-fit on the pulp temperature for each segment was performed with MATLAB’s fit() function under a custom exponential decay model of the form

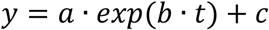

in which the time constant was defined as

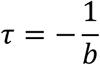

##### Threshold response temperature

Threshold response temperature calculation was performed in MATLAB. Intradental neuronal response temperatures were consolidated, sorted from high to low, and plotted cumulatively. The slope of the curve represented the neuronal response density (unit: number of responses/°C). To capture the global neuronal response density transition, a piecewise (up to 10 segments) linear curve fit was performed, and the angles between adjacent segments (0∼90°; higher values indicate a steeper transition) were calculated at the critical points to quantify the slope transition. The threshold response temperature was determined at the critical point with the highest response density transition.

*AUC*

Area Under the Curve measurements were performed in MATLAB. All subzero values in normalized GCaMP6f fluorescence traces (induced by insufficient subtraction of adjacent cell fluorescence in the neuropil region) were substituted with zeros. The traces were then normalized to the range [0, 1], and AUC was calculated for the designated paradigm using MATLAB’s trapezoidal numerical integration function trapz().

##### Maximum normalized GCaMP6f fluorescence intensity

Maximum normalized GCaMP6f fluorescence intensity was calculated as the responder’s maximum intensity among its response instances.

#### Retrograde labeling of intradental HTMRs

Retrograde labeling of intradental neurons via cholera toxin B-subunit (CTB) was performed in adherence to a published method(*30*, *35*). Briefly, CTB 647 was diluted to 100ug/10ul in DDwater. Adult mice (>6 weeks of age) were anesthetized with isoflurane (4% for induction;1.5∼2% for maintenance), ophthalmic ointment was applied to both eyes, and body temperature was maintained with a hand warmer. Oral cavity access was achieved via a custom-designed stage, on which mice’s maxillary and mandibular incisors were loaded in counter directions to open the mouth, and cheeks and tongue were retracted laterally using surgical retractors to expose the molars. Shallow occlusal dentin exposure was performed with a ¼ round carbide dental bur (HSB-401152, House Brand) attached to a micromotor handpiece (35,000RPM). OptiBond S (Kerr), a hydrophobic bonding agent, was applied to cover soft tissues surrounding the molar to prevent nonspecific labeling. CTB volume (∼1μL per tooth) was delivered in 3 applications over 15 minutes, allowing each application to gently dry. The exposed dentin was then sealed with dental filling (Flow-It ALC B2; Pentron) and UV-cured. Mice were allowed to recover for 16 hours before TG harvesting for in situ hybridization.

##### Pulpitis model

Induction of acute pulpitis via application of inflammatory soup was achieved using a surgical protocol similar to that described above for retrograde labeling of intradental HTMRs. Briefly, following surgical exposure of dentin, 2µL of inflammatory soup (using an established recipe(*51*, *52*) consisting of a mixture of inflammatory mediators consisting of bradykinin (1mM), serotonin (1mM), histamine (1mM), and prostaglandin E2 (100µM) in 1x PBS, was applied to the exposed dentin in multiple small-volume applications. Care was taken to ensure each application cast a thin layer of liquid on the occlusal surface without spillage to surrounding tissues. A gentle vacuum was used to promote convection near the molars and facilitate drug absorption. After full dispensing of 2µL, the exposed dentin was sealed with a dental filling (Flow-It ALC B2; Pentron) and UV-cured. Mice were returned to their home cages and allowed to recover for 48 hours before in vivo TG imaging or tissue harvesting for pulpal immune cell IHC quantification. Littermate controls either underwent this procedure but instead received PBS or underwent no surgical procedure.

#### In situ hybridization (ISH) of TG in tissue sections

TG ISH was performed in fresh frozen samples harvested from mice following immediate euthanasia with CO2. Dissected TG were immediately embedded in OCT medium and flash frozen on dry ice for cryosectioning (Leica CM1950), sectioned at a thickness of 20μm onto Superfrost Plus Slides (Fisher #12-550-15), and slides were stored in −80°C prior to use. In situ hybridization (ISH) was performed following an adaptation of hybridization chain reaction (HCR) protocol version 3(*14*, *30*, *35*, *72*).

HCR was performed using buffers, hairpin amplifiers, and probes against transcripts for mouse genes *Scn10a* (PRA454), *S100b* (PRA454), *Trpv1* (PRB721), *Trpa1* (PRF447), and *Trpm8* (PRB982), and *GCaMP* (RTE507) from Molecular Instruments. Briefly, slides were first fixed on ice with 4% PFA/1X PBS for 15 minutes, followed by three washes with 1X PBS. Acetylation was performed using Milli-Q water in 0.3% v/v acetic anhydride and 0.1M triethanolamine. After three additional washes in 1X PBS, the slides were dehydrated via an ethanol series in 50%, 70%, 100%, and fresh 100% ethanol for 5 minutes at each concentration respectively, followed by three rinses in 2X saline sodium citrate (SSC). Then, slides were prehybridized in a Coverwell chamber (Grace Biolabs) with hybridization buffer at 37°C for 10 minutes in a humidified hybridization oven. Meanwhile, a working hybridization solution was prepared by adding probes with adapters at a concentration of 4nM per probe to the pre-warmed hybridization buffer at 37°C. The prehybridization buffer was then replaced with this working hybridization solution, allowing the sections to be incubated at 37°C for 1-3 days in a humidified hybridization oven. The next day, hybridized slides were washed in a 100% wash buffer for 5 minutes at 37°C. This is followed by a graded series of 10 minutes washes at 37°C consisting of 75%, 50%, 25% wash buffer diluted in 5X SSC with 0.1% Tween 20 (SSCT), and finally a 100% 5X SSCT for 5 minutes at room temperature (RT). The slides were pre-incubated for 30 minutes with an amplification buffer, which was followed by overnight incubation in a working amplification solution at RT in a dark, humidified chamber. The working amplification solution containing fluorophore-conjugated B1-B6 amplifier hairpins were made freshly before use in accordance with the manufacturer recommendations and diluted using the amplification buffer. Fluorophores (Alexa 546, Alexa 594, Alexa 750) were selected to avoid overlaps with endogenous fluorescence present in the samples. After amplification, the slides were washed two times with 5X SSCT for 15 minutes with gentle agitation. Sections were then mounted in Imaging Buffer (3U/mL pyranose oxidase, 0.8% D-glucose, 2X SSC, 10mM Tric HCl pH 7.4, and 400U RNAse inhibitor), coverslipped, and sealed with Cytobond (Scigene) before imaged using confocal microscopy (Olympus FV3000, Evident Scientific, Inc.). As multiple rounds of multi-plexed ISH were required to complete imaging of all probes, following imaging of the first round of probes, slides were washed in 2X SSC followed by stripping of DNA probes and amplifiers via digestion with 250 U/ml RNase-free DNase (Roche Diagnostics). The hybridization procedure was repeated with additional probes for a total of 3 rounds as described previously. Scoring for probe expression overlap with CTB or the genetically-encoded Cre reporter was performed by drawing ROIs in ImageJ in cells of interest followed by manual scoring for probe expression based on the presence of visible signal within the ROI above surrounding background fluorescence. To facilitate manual scoring of high-ROI-count images, a custom MATLAB application was adopted to extract and display individual ROI fluorescence overlays, ensuring a trackable workflow.

#### TG terminal ending visualization and immunohistochemistry for pulpitis validation

##### AAV Viral Delivery

For labeling of *Bmpr1b-*Cre somatosensory neuron terminals, postnatal day 1-3 *Bmpr1b-*Cre pups received intraperitoneal AAV injection of AAV9-Cre. Mouse pups were briefly anesthetized prior to i.p. injection by moving them to a plastic petri dish on ice. Pups were injected using a Hamilton syringe (Hamilton Company, Reno, USA) containing 10uL volume of 10^12^ viral genomes of ssAAV-9/2-CAG-dlox-tdTomato(rev)-dlox-WPRE-bGHp(A) (Physical titer: 5.5 x 10^12^ vg/ml, catalog # v167-9, University of Zürich Viral Vector Facility) diluted in sterile saline. Tissues of interest were collected from animals at adulthood (8-12 weeks) following perfusion, decalcification, and sample preparation as described below.

##### Perfusion and decalcification

To prepare tissue samples for histology or terminal ending visualization, adult (8-12 weeks) animals were euthanized via CO2, perfused with 30mL of 4°C 1X PBS followed by 4% PFA/1X PBS. Tissues of interest (skulls or mandibles) were dissected and post-fixed overnight in 4% PFA/1X PBS, after which skull samples were hemisectioned. Samples were then decalcified in 0.1M EDTA/1X PBS at 42°C shaking (350RPM) for 6 (skulls) or 7 (mandibles) days, with buffer exchange every 2 days.

##### Whole-mount terminal ending visualization of lip tissue

To visualize *Bmpr1b*-Cre terminal endings within the lip, whole-mount clearing was performed on the decalcified AAV injected *Bmpr1b*-Cre skull samples with a modified KneEZ Clear protocol(*75*). Samples underwent a series of graded tetrahydrofuran dehydration and rehydration followed by equilibration in EZ Clear solution for refractive index matching. Endogenous TdT fluorescence within the lip tissue was imaged using an Olympus FV3000 confocal microscope under stitched Z-stacks.

##### Cryosectioning

Cryosectioning was performed on decalcified hemi-sectioned skulls or mandibles prior to examining molar endogenous TdT fluorescence or performing immunohistochemistry. Samples were cryoprotected in 30% sucrose/1X PBS for 48 hours until sunk, embedded in OCT, flash-frozen with dry ice, and stored at −80°C until sectioning. OCT-embedded tissues were sectioned (Leica CM1950) at 20μm (molars) or 50μm (hemisected whole skulls) and thaw-captured onto Gelatin Subbed Slides (Southern Biotech, #SLD01-BX) or Superfrost Plus Slides (Fisher Scientific). Slides were stored at −80°C until endogenous TdT fluorescence was imaged or immunohistochemistry was performed.

##### Visualization of intradental innervation (endogenous TdT fluorescence)

To visualize *Bmpr1b*-Cre terminal endings within the molar, slides with skull sections were thawed at room temperature and endogenous TdT fluorescence was imaged using an Olympus FV3000 confocal microscope under stitched Z-stacks.

##### Pulpal immune cell validation (immunohistochemistry)

Immunohistochemistry (IHC) was performed on slides with mandible sections to examine the presence of pulpal immune cells as described previously(*30*). Briefly, slides were first thawed at room temperature. The tissue area on each slide was outlined with an ImmEdge hydrophobic barrier pen (Vector Laboratories). Tissue sections were post-fixed using 1X PBS/4% PFA for 10 minutes following three washes with 1X PBS. Nonspecific binding was blocked by incubating the sections for 1 hour at room temperature in 10% Normal Goat or Donkey Serum containing 0.1% Triton X-100/1X PBS. Sections were subsequently incubated with primary antibody overnight at 4°C, followed by three washes in 1X PBS and incubation in secondary antibody for 2 hours at RT while protected from light. After three final washes with 0.1% Triton X-100/1X PBS, the sections were mounted in Prolong Diamond, either with or without DAPI (Life Technologies), coverslipped, and imaged via confocal microscopy (Olympus FV3000, Evident Scientific, Inc.).

##### Quantification of immune cell fluorescence intensity

Fluorescence quantification of pulpal CD45 was performed using a custom MATLAB code. Briefly, image stacks consisting of a DAPI channel and an immune cell channel were inputted, and the pulp ROI was defined by thresholding the DAPI channel, followed by binary operations to remove small objects and fill holes and a visual examination for accuracy. Pixel values of the immune cell channel inside the pulp ROI were then extracted, sorted from low to high, and fitted with a 2-component Gaussian Mixture distribution Model (GMM) using MATLAB’s fitgmdist() function. The GMM’s low-intensity component represented fluorescence intensity contribution from native pulpal immune cells, while the high-intensity component accounted for localized immune cell enrichments. This approach enabled unbiased quantification of pulpal immune cell fluorescence intensity across different experimental conditions.

#### Quantification and Statistical Analysis

Statistical analyses were performed in GraphPad prism. Welch’s unpaired t test was adopted to compare two groups, while ordinary one-way ANOVA was performed for comparisons between more than two conditions. Statistical significance was defined as p value <0.05. For each analysis, relevant information including statistical methods, error bars, p values, and sample sizes were described in the corresponding figure legends.

Figs. S1 to S3

---

**Fig. S1:**
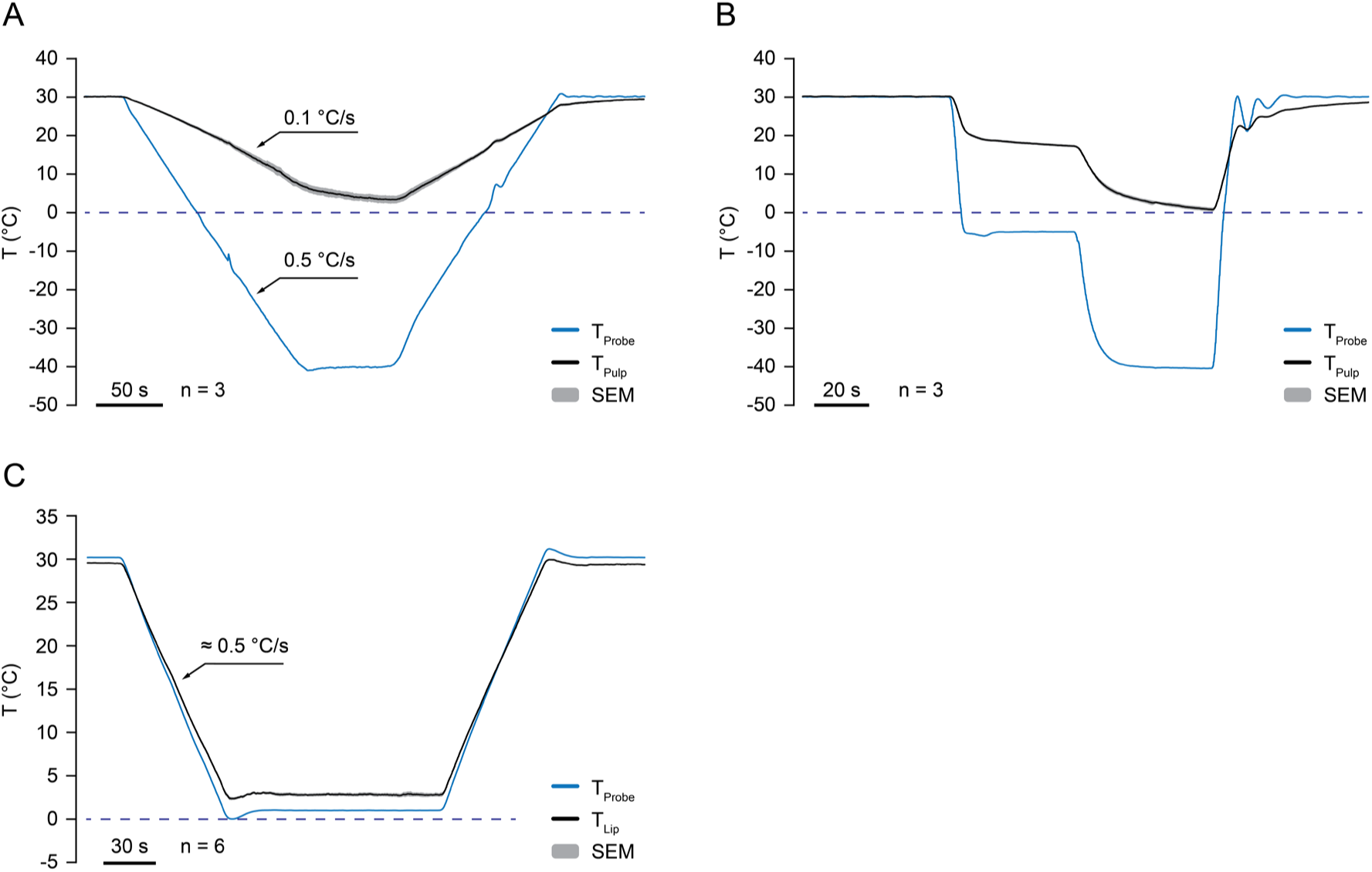
Probe-tissue temperature correlations. Related to Fig. 2, 3, and 4. **(A)** Graph showing the probe temperature (blue) and corresponding tooth pulp temperature (black) during slow molar cooling. Traces show the average; shading indicates SEM. n = 3. Scale bar, 50s. **(B)** Graph showing the probe temperature (blue) and corresponding tooth pulp temperature (black) during stepwise fast molar cooling. Traces show the average; shading indicates SEM. n = 3. Scale bar, 20s. **(C)** Graph showing the probe temperature (blue) and corresponding lip surface temperature (black) during slow lip cooling. Traces show the average; shading indicates SEM. n = 6. Scale bar, 30s.

**Fig. S2.**
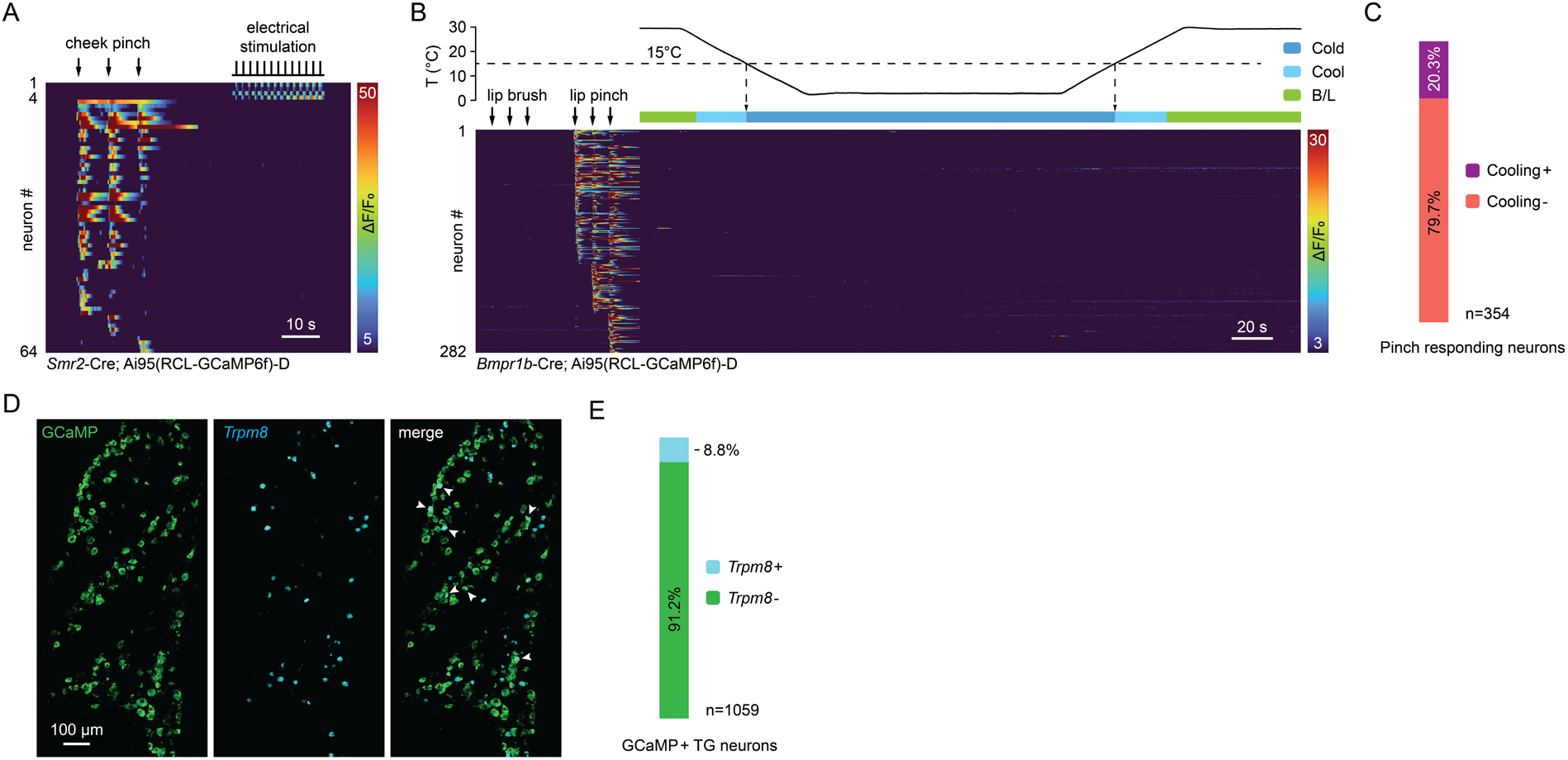
Smr2-Cre does not capture intradental electrical responders; functional and molecular validation of Bmpr1b-Cre strain. Related to. Fig. 4**. (A)** Heatmap showing in vivo GCaMP responses from TG neurons to cheek pinch and electrical stimulation to mandibular m1. TG neurons respond robustly to cheek pinch but rarely to m1 electrical stimulation. Data are obtained from *Smr2*-cre;GCaMP6f mice. n = 60 cheek pinch responding neurons and n = 4 tooth electrical responding neurons across 3 TGs. Scale bar, 10s. **(B)** Heatmap showing in vivo GCaMP responses of *Bmpr1b*-Cre TG neurons responding to lip pinch but neither to slow lip cooling nor to lip brush. n = 282 neurons across 6 TGs. Scale bar, 20s. **(C)** Slice plot showing the percentage of *Bmpr1b*-Cre lip pinch responding neurons that were responsive or unresponsive to slow lip cooling. n = 354 pinch responding neurons across 6 TGs. (D and E) *Bmpr1b*-Cre-labeled TG neurons minimally express *Trpm8*. (D) Representative image of ISH staining for GCaMP (green) and *Trpm8* (cyan). Scale bar, 100µm. (E) Slice plot showing the percentage of *Bmpr1b*-Cre-labeled TG neurons that express or do not express *Trpm*8. n = 1059 GCamp+ neurons in 9 sections from 3 TGs

**Fig. S3.**
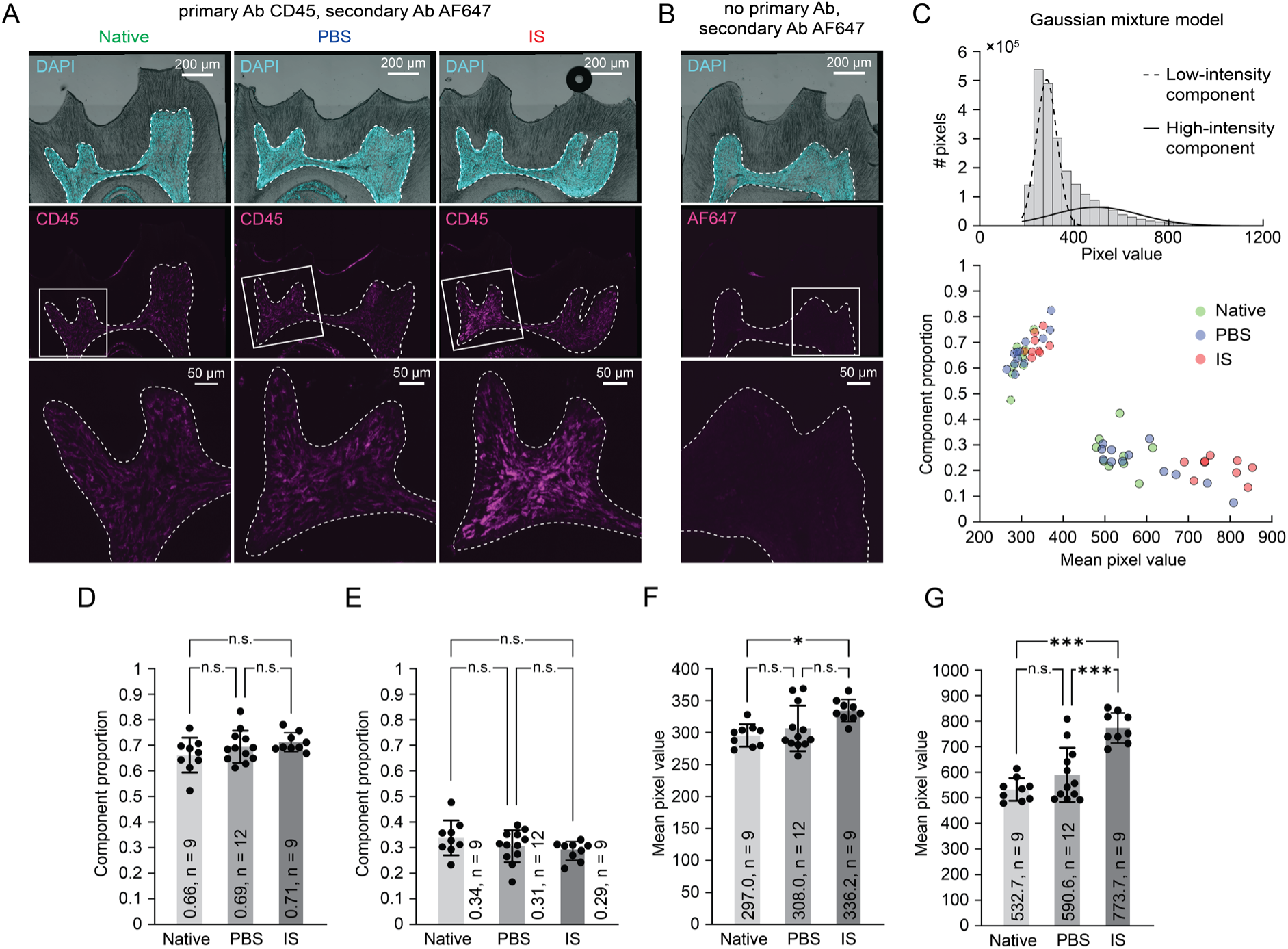
Immunohistochemistry of immune cells in dental pulp. Related to. Fig. 5 **(A)** Representative image of CD45 immunohistochemistry in mandibular first molars receiving no surgical procedure (left column), PBS treatment (middle column), or IS treatment (right column). Top row: Merged image overlaid with DAPI staining labeling nuclei of pulp cells and brightfield image of mandibular m1 cross section. Enamel has been fully removed during demineralization. Dashed lines indicate the pulp-dentin border. Scale bar, 200µm. Middle row: Immunostaining for CD45. Bottom row: Magnified image of the white rectangle. Scale bar, 50µm. This experiment was repeated in n = 9 (native tooth, IS) or n = 12 (PBS) sections from 3 (native tooth, IS) or 4 (PBS) mandibular m1s. **(B)** Representative image of immunostaining of demineralized mandibular m1 sections in the absence of the CD45 primary antibody. Top panel: Merged image overlaid with DAPI staining labeling nuclei of pulp cells and brightfield image of mandibular m1 cross section. Dashed lines indicate the pulp-dentin border. Scale bar, 200µm. Middle panel: Immunostaining exhibits no AF647+ signal with the CD45 primary antibody omitted. Bottom panel: Magnified image of the white rectangle. Scale bar, 50µm. This experiment was repeated in n = 3 (native tooth, IS) or n = 4 (PBS) sections from 3 (native tooth, IS) or 4 (PBS) mandibular m1s. **(C)** 2-component Gaussian Mixture distribution Model (GMM) analysis of pulpal CD45 fluorescence intensity. Top: Representative graph showing the GMM fitted to pulpal pixel values. Histogram shows the raw pixel values. Dashed trace indicates the low-intensity Gaussian component; solid trace indicates the high-intensity Gaussian component. Bottom: Graph showing the GMM low- and high-intensity component distributions over their mixing proportions and mean values for native tooth (green), PBS (blue), and IS (red) conditions. Dashed datapoint borders indicate low-intensity components and solid borders indicate high-intensity components. Results in (C) are from primary data represented by (A). n = 9 (native tooth, IS) or n = 12 (PBS) sections from 3 (native tooth, IS) or 4 (PBS) mandibular m1s. **(D and E)** Bar graphs showing the mixing proportions of GMM low-intensity components (D) and high-intensity components (E). Plotted individual data points represent the components’ mixing proportions in each condition. Bar shows the mean value and error bars indicate the SEM. Results in (D) and (E) are from primary data represented by (A). n = 9 (native tooth, IS) or n = 12 (PBS) sections from 3 (native tooth, IS) or 4 (PBS) mandibular m1s. For both low- and high-intensity components, p = 0.7676 (IS vs. PBS), p = 0.1724 (IS vs. native tooth), and p = 0.4168 (PBS vs. native tooth) using one-way ANOVA with Tukey’s correction. **(F and G)** Bar graphs showing the mean pixel values of GMM low-intensity components (F) and high-intensity components (G). Plotted individual data points represent the components’ mean pixel values in each condition. Bar shows the mean value and error bars indicate the SEM. Results in (F) and (G) are from primary data represented by (A). n = 9 (native tooth, IS) or n = 12 (PBS) sections from 3 (native tooth, IS) or 4 (PBS) mandibular m1s. For low-intensity components, p = 0.0582 (IS vs. PBS), p = 0.0111 (IS vs. native tooth), and p = 0.6184 (PBS vs. native tooth) using one-way ANOVA with Tukey’s correction. For high-intensity components, p < 0.0001 (IS vs. PBS), p < 0.0001 (IS vs. native tooth), and p = 0.2368 (PBS vs. native tooth) using one-way ANOVA with Tukey’s correction.

